# Fibrosis protein-protein interactions from Google matrix analysis of MetaCore network

**DOI:** 10.1101/2021.10.20.465138

**Authors:** Ekaterina Kotelnikova, Klaus M. Frahm, Dima L. Shepelyansky, Oksana Kunduzova

## Abstract

Protein-protein interactions is a longstanding challenge in cardiac remodeling processes and heart failure. Here we use the MetaCore network and the Google matrix algorithms for prediction of protein-protein interactions dictating cardiac fibrosis, a primary causes of end-stage heart failure. The developed algorithms allow to identify interactions between key proteins and predict new actors orchestrating fibroblast activation linked to fibrosis in mouse and human tissues. These data hold great promise for uncovering new therapeutic targets to limit myocardial fibrosis.

## 1. Introduction

Cardiovascular disease, a class of diseases that impact cardiovascular system, is responsible for 31% of all deaths and remains the leading cause of mortality worldwide [1]. Myocardial fibrosis is a central element of cardiac remodeling that leads to human failure and death [2]. Myocardial fibrosis results from uncontrolled fibroblast activity and excessive extracellular matrix deposition [2]. Although a number of factors have been implicated in orchestrating the fibrotic response, tissue fibrosis is dominated by a central mediator: transforming growth factor-*β* (TGF-*β*) [3]. Sustained TGF-*β* production leads to a continuous cycle of growth factor signaling and dysregulated matrix turnover. However, despite intensive research, the factors that orchestrate fibrosis are still poorly understood and as a result, effective strategies for reversing fibrosis are lacking [2,4]. Considering the complex heterogeneity of fibrosis, research strategy on a system-level understanding of the disease using mathematical modeling approaches is a driving force to dissect the complex processes involved in fibrotic disorders. Recently, we have reproduced the classic hallmarks of aberrant cardiac fibroblast activation leading to fibrosis including high collagen production and deposition [5]. Tools designed in this work, with RNA sequence data sets, enable analyses to help generate hypotheses about a gene’s function in activated fibroblasts.

The present work is based on the recent clinical results presented in [5] which allowed to determine the protein response detected in fibrosis tissue of mouse and human as a feedback on TGF protein stimulation, which is known to play an important role [3]. These experiments allowed to determine proteins with most positive and most negative response. We use the proteins with top 20 positive and top 20 negative strongest response. Their names are given in Table 1 marked by indexes *K_u_* = 1, 2, …, 20; *K_d_* = 1, 2, …, 20. These proteins are ordered monotonically from the strongest *K_u_* = 1 to to weakest *K_u_* = 20 positive responses; the same monotonic ordering is done by modulus of negative response with strongest *K_d_* = 1 to weakest *K_d_* = 20 responses. In addition there are 4 proteins TGF-*β* with indexes *K_t_* = 1, 2, 3, 4 used in experiments [5]. These 44 proteins form the internal selected fibrosis group. For the analysis of protein-protein interactions (PPI) characterizing fibrosis we add a group of 10 external proteins with indexes as *K_x_* = 1, 2, …, 10 (their choice will be explained below). Thus in total we have the PPI fibrosis network with 54 proteins (nodes). They are ordered by their global index *K_g_* = 1, 2, …, 54 as it is shown in Table 1 (first 4 *K_t_*, then 20 *K_u_*, 20 *K_d_* and 10 *K_x_*).

**Table 1.** Table of the subset of *N_r_* = 54 selected fibrosis proteins (nodes). Here *K_g_* represents the global index of this group, *K_t,u,d,x_* represent the index of the four subgroups of 4 TFG-*β* proteins, 20 up-proteins, 20 down-proteins and 10 additional X-proteins; *K*(*K*\*) represents the local PageRank (CheiRank) index inside this group and 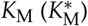 represents the PageRank (CheiRank) index for the global MetaCore network of *N* = 40079 nodes; the last column gives the associated protein names.

| $K_g$ | $K_{t,u,d,x}$ | $K$ | $K^*$ | $K_M$ | $K_M^*$ | Protein |
| --- | --- | --- | --- | --- | --- | --- |
| 1 | $K_t = 1$ | 30 | 37 | 10780 | 26299 | TGF-β 0 |
| 2 | $K_t = 2$ | 9 | 14 | 235 | 5690 | TGF-β 1 |
| 3 | $K_t = 3$ | 13 | 33 | 968 | 25073 | TGF-β 2 |
| 4 | $K_t = 4$ | 20 | 45 | 4726 | 29508 | TGF-β 3 |
| 5 | $K_u = 1$ | 46 | 35 | 28737 | 25928 | ADAMTS16 |
| 6 | $K_u = 2$ | 17 | 34 | 3478 | 25137 | FGF21 |
| 7 | $K_u = 3$ | 52 | 39 | 40048 | 28152 | TNFSF18 |
| 8 | $K_u = 4$ | 16 | 26 | 2467 | 19160 | ACAN |
| 9 | $K_u = 5$ | 14 | 31 | 1489 | 24511 | RPH3A |
| 10 | $K_u = 6$ | 42 | 46 | 26600 | 29559 | ADAMTS8 |
| 11 | $K_u = 7$ | 51 | 47 | 34769 | 39960 | MEGF6 |
| 12 | $K_u = 8$ | 40 | 38 | 26295 | 27326 | SV2B |
| 13 | $K_u = 9$ | 44 | 48 | 27111 | 36021 | C1QTNF3 |
| 14 | $K_u = 10$ | 50 | 49 | 34616 | 39841 | ANO4 |
| 15 | $K_u = 11$ | 32 | 24 | 12696 | 16566 | IL11 |
| 16 | $K_u = 12$ | 43 | 30 | 26624 | 23640 | CDH10 |
| 17 | $K_u = 13$ | 26 | 50 | 7263 | 30243 | HTR2B |
| 18 | $K_u = 14$ | 19 | 16 | 4647 | 6551 | LAMA1 |
| 19 | $K_u = 15$ | 28 | 36 | 8342 | 26295 | LAMA1 |
| 20 | $K_u = 16$ | 18 | 17 | 4021 | 8252 | RAPGEF4 |
| 21 | $K_u = 17$ | 48 | 51 | 29945 | 36964 | DNER |
| 22 | $K_u = 18$ | 36 | 18 | 22159 | 8569 | GALNT3 |
| 23 | $K_u = 19$ | 47 | 23 | 29145 | 15531 | ACSBG1 |
| 24 | $K_u = 20$ | 37 | 20 | 24786 | 8735 | OLFM2 |
| 25 | $K_d = 1$ | 35 | 40 | 19039 | 28262 | CLEC3B |
| 26 | $K_d = 2$ | 41 | 41 | 26477 | 28290 | SCARA5 |
| 27 | $K_d = 3$ | 39 | 22 | 26109 | 11185 | SLC10A6 |
| 28 | $K_d = 4$ | 24 | 44 | 6360 | 29204 | CXCL5 |
| 29 | $K_d = 5$ | 33 | 19 | 14952 | 8729 | MYOC |
| 30 | $K_d = 6$ | 22 | 28 | 5961 | 22288 | IFITM1 |
| 31 | $K_d = 7$ | 21 | 13 | 5599 | 4483 | ANGPTL4 |
| 32 | $K_d = 8$ | 38 | 25 | 25538 | 17434 | SELENBP1 |
| 33 | $K_d = 9$ | 34 | 52 | 18938 | 33179 | FMO1 |
| 34 | $K_d = 10$ | 49 | 53 | 34080 | 39427 | GPR88 |
| 35 | $K_d = 11$ | 23 | 27 | 6276 | 22141 | HMGCS2 |
| 36 | $K_d = 12$ | 53 | 43 | 37060 | 28328 | LGI2 |
| 37 | $K_d = 13$ | 29 | 11 | 9162 | 2485 | PTN |
| 38 | $K_d = 14$ | 11 | 15 | 513 | 5974 | ADORA2A |
| 39 | $K_d = 15$ | 27 | 29 | 7789 | 22652 | GFRA1 |
| 40 | $K_d = 16$ | 25 | 21 | 6718 | 8844 | IL1R2 |
| 41 | $K_d = 17$ | 54 | 42 | 35446 | 28306 | IL1R2 |
| 42 | $K_d = 18$ | 31 | 12 | 12148 | 3444 | PEG10 |
| 43 | $K_d = 19$ | 45 | 54 | 27829 | 36195 | FMO2 |
| 44 | $K_d = 20$ | 15 | 32 | 1973 | 24994 | COX4I2 |
| 45 | $K_x = 1$ | 1 | 4 | 3 | 13 | β-catenin |
| 46 | $K_x = 2$ | 2 | 1 | 4 | 6 | p53 |
| 47 | $K_x = 3$ | 3 | 2 | 11 | 10 | ESR1 |
| 48 | $K_x = 4$ | 4 | 5 | 13 | 25 | STAT3 |
| 49 | $K_x = 5$ | 5 | 3 | 22 | 11 | RelA |
| 50 | $K_x = 6$ | 6 | 6 | 38 | 82 | PPAR-γ |
| 51 | $K_x = 7$ | 7 | 8 | 111 | 767 | IKK-β |
| 52 | $K_x = 8$ | 8 | 7 | 179 | 198 | SNAIL1 |
| 53 | $K_x = 9$ | 10 | 9 | 237 | 1520 | MMP-14 |
| 54 | $K_x = 10$ | 12 | 10 | 578 | 2123 | Flotillin-1 |

To analyze the properties of this PPI fibrosis network we use the developed commercial MetaCore network database of Clarivate [6]. This network database has been shown to be useful for analysis of various specific biological problems (see e.g. [7,8]). At present the MetaCore network has *N* = 40079 nodes with *N*_ℓ_ = 292191 links (without self connections) with on average *n*_ℓ_ = *N*_ℓ_ / *N* ≈ 7.3 links per node [9]. The nodes are given mainly by proteins but also there are certain molecules and molecular clusters catalyzing the interactions with proteins. This MetaCore PPI network is directed and non-weighted. Also its network links mark the bi-functional nature of interactions leading to the activation or the inhibition of one protein by another one. For some nodes link action is neutral or unknown.

For the investigation of fibrosis PPI network we use the Google matrix algorithms developed for the analysis of the World Wide Web [10,11] and other directed networks like Wikipedia networks, world trade networks and others (see review [12]). Such an approach to network characterization is based on the concept of Markov chains invented by Markov in an article published 1906 in the proceeding of the Kazan University [13].

The important method for analysis of directed networks is the reduced Google matrix (REGOMAX) algorithm developed and described in detail in [14,15]. The REGOMAX algorithm has been applied to PPI networks of SIGNOR database as reported in [16,17]. However, the number of nodes in the SIGNOR database is by a factor ten smaller than the number of nodes of the MetaCore network and its use can only be considered as a test bed for of the numerical algorithms and its conceptional base. A first description of the statistical properties of the global MetaCore network, including PageRank, CheiRank and REGOMAX characteristics, was presented in [9]. However, this work only represents a statistical study of the MetaCore network without any applications to a concrete biological problem. In this work we apply the REGOMAX analysis to the specific biological problem of fibrosis.

The important feature of the REGOMAX algorithm is that it constructs the Google matrix of a selected subset of nodes *N_r_* ≪ *N* (here we have *N_r_* = 54) taking into account not only direct links between these *N_r_* nodes but also all indirect pathways connecting them via the global MetaCore network of much larger size *N*. The efficiency the REGOMAX approach was demonstrated for various applications concerning the Wikipedia and world trade networks [18–21] and we also expect that this method will provide useful and new insights in the context of fibrosis protein-protein interactions using the MetaCore network.

The paper constructing as follows: Section 2 describes the data sets and Google matrix algorithms, Section 3 presents the obtained results of the reduced Google matrix and sensitivity analysis for the particular group of 54 proteins (of Table 1) we consider here and Section 4 provides the discussion of the results and the conclusion; In the Appendix we provide additional Figures and a simple analytical estimate for the sensitivity matrix to which we refer in the main part of the work; more detail numerical data files obtained from the Google matrix computations are available at [22].

## 2. Data sets and methods

Here we describe the data sets, construction rules and algorithms of Google matrix.

### 2.1. Network data sets

The global MetaCore PPI network contains *N* = 40079 nodes with *N*_ℓ_ = 292191 links (without self connections). The number of activation/inhibition links is *N*_ℓ+_ / *N*_ℓ−_ = 65157/49321 ≃ 1.3 and the number of neutral links is *N*_ℓ*n*_ = *N* – *N*_ℓ+_ – *N*_ℓ−_ = 177713. Here we mainly present the results without taking into account the bi-functional nature of links. However, a part of the results takes into account this bi-functionality of links using the Ising Google matrix approach described in [9,17]. The subset of selected *N_r_* = 54 fibrosis proteins (nodes) is given in Table 1; these nodes are represented by 4 TGF-*β* proteins/nodes (*K_t_* = 1, 2, 3, 4), 20 “up-proteins” (*K_u_* = 1, …, 20), 20 “down-proteins” (*K_d_* = 1, …, 20), both obtained from experiments [5] (as described above) and 10 new “X-proteins” (or “X-nodes”; *K_x_* = 1, …, 10) whose selection is explained later. The TGF-*β* 4 nodes correspond to different isoforms of this protein.

The Google matrix approach used in this work is explained in detail in [10–12] and the related REGONAX algorithm is described in [9,14,15,17]. Below we present a short description of these methods following mainly the presentation given in [9], keeping the same notations.

### 2.2. Google matrix construction, PageRank and CheiRank

First, we construct the Google matrix *G* of the MetaCore network for the simple case where the bi-functional nature of links is neglected. Furthermore, the directed links are non weighted. First one defines an adjacency matrix with elements *A_ij_* being equal to 1 if node *j* points to node *i* and equal to 0 otherwise. In the next step, the stochastic matrix *S* describing the node-to-node Markov transitions is obtained by normalizing each column sum of the matrix *A* elements to unity. For dangling nodes *j* corresponding to zero columns of *A*, i.e. *A_ij_* = 0 for all nodes *i*, the corresponding elements of *S* are defined by *S_ij_* = 1/ *N*. The stochastic matrix *S* describes a Markov process on the network: a random surfer jumps from node *j* to node *i* with the probability *S_ij_*, therefore following the directed links. The column sum normalization ∑_*i*_ *S_ij_* = 1 ensures the conservation of probability. The elements of the Google matrix *G* are then defined by the standard form

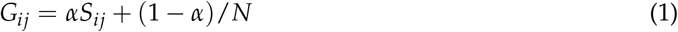

where *α* = 0.85 is the usual damping factor [10,11]. The Google matrix is also column sum normalized and now the random surfer jumps on the network in accordance to the stochastic matrix *S* with a probability *α* and with a complementary probability (1 – *α*), to an arbitrary random node of the network. The damping factor allows to escape from possible isolated communities and ensures that the Markov process converges for long times rather quickly to a uniform stationary probability distribution. The latter is given by the PageRank vector *P* which is the right eigenvector of the Google matrix *G* corresponding to the leading eigenvalue, here *λ* = 1. The corresponding eigenvalue equation is then *GP* = *P*. According to the Perron-Frobenius theorem, the PageRank vector *P* has positive elements and their sum is normalized to unity. The PageRank vector elements *P*(*j*) gives the probability to find the random surfer on the node *j* at the stationary state of the Markov process. Thus, all the nodes can be ranked by monotonically by decreasing PageRank probability. The PageRank index *K*(*j*) gives the rank of the node *j* with the highest (lowest) PageRank probability *P*(*j*) corresponding to *K*(*j*) = 1 (*K*(*j*) = *N*). The PageRank probability *P*(*j*) is proportional, on average, to the number of ingoing links pointing to node *j*. However, it also takes into account the “importance” (i.e. PageRank probability) of the nodes having a direct link to *j*.

It is also useful to consider a network obtained by the inversion of all link directions. For this inverted network, the corresponding Google matrix is denoted *G*\* and the corresponding PageRank vector, called the CheiRank vector *P*\*, is defined such as *G*\**P*\* = *P*\*. A detailed statistical analysis of the CheiRank vector can be found in [23,24] (see also [12]). Similarly to the PageRank vector, the CheiRank probability *P*\* (*j*) is proportional, on average, to the number of outgoing links going out from node *j*. The CheiRank index *K*\*(*j*) is also defined as the rank of the node *j* according to decreasing values of the CheiRank probability *P*\*(*j*).

### 2.3. Reduced Google matrix

The concept of the REGOMAX algorithm was introduced in [14] and a detailed description of the first applications to groups of political leaders having articles in Wikipedia networks (different language editions) can be found in [15]. This algorithm determines effective interactions between a selected subset of *N*_r_ nodes enclosed in a global network of size *N* ≫ *N*_r_. These interactions are determined taking into account direct and all indirect transitions between *N*_r_ nodes via all the other *N*_s_ = *N* – *N*_r_ nodes of the global network. We note that quite often in certain network analyses only direct links of a subset of elected *N*_r_ nodes are taking into account and their indirect interactions via the global network are omitted thus clearly missing the important interactions.

On a mathematical level the REGOMAX approach uses ideas similar to those of the Schur complement in linear algebra (see e.g. [25]) and quantum chaotic scattering in the field of quantum chaos and mesoscopic physics (see e.g. [26,27]). The Schur complement was introduced by Issai Schur in 1917 (see history in [25]). In the context of Markov chains this approach was discussed in [28]. It is clear that the Schur complement attracted a lot of studies from far 1917. However, there are new elements, developed in [14,15], related to a specific matrix decomposition of the Schur complement which allows to understand its new features and to compute efficiently (numerically) the three related matrix components in the framework of the reduced Google matrix approach for very large networks (e.g. *N* ~ 5 × 10^6^ as for English Wikipedia).

We write the full Google matrix *G* of the global network in the block form

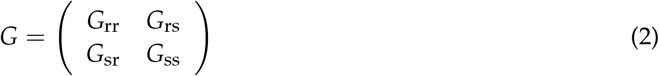

where the label “r” refers to the nodes of the reduced network, i.e. the subset of *N*_r_ nodes, and “s” to the other *N*_s_ = *N* – *N*_r_ nodes which form the complementary network acting as an effective “scattering network”. The reduced Google matrix *G*_R_ acts on the subset of *N*_r_ nodes, has the size *N*_r_ × *N*_r_. It is defined by

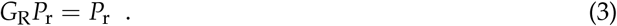

Here *P*_r_ is a vector of size *N*_r_, its components are the normalized PageRank probabilities of the *N*_r_ nodes, 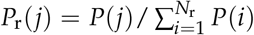. The REGOMAX approach allows to find an effective Google matrix for the subset of *N*_r_ nodes keeping fixed the relative ranking probabilities between these nodes. The reduced Google matrix *G*_R_ has the form [14,15]

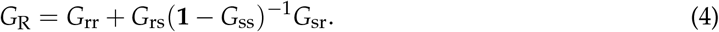

Furthermore it satisfies the relation (3) and it is also column sum normalized. The reduced Google matrix *G*_R_ can be represented as the sum of three components [14,15]:

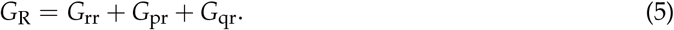

Here, the first component, *G*_rr_, corresponds to the direct transitions between the *N*_r_ nodes; the second component, *G*_pr_, is a matrix of rank 1 with all the columns being proportional (actually approximately equal to the reduced PageRank vector *P*_r_); the third component, *G*_qr_, describes all the “interesting indirect pathways” passing through the global network of *G* matrix. Without going into the details we mention here that mathematically (and also numerically) *G*_pr_ is obtained from (4) by extracting the contribution of the leading eigenvector of *G*_ss_ (which is very close to the PageRank of the complementary scattering network of *N*_s_ nodes) whose eigenvalue is close to unity but it is *not exactly* unity since *G*_ss_ is not column normalized and there is small escape probability from the *N*_s_ scattering nodes to the selected subset with *N*_r_ nodes. This eigenvector dominates therefore the matrix inverse in (4) and its contribution produces the rank 1 matrix *G*_pr_ and the remaining contributions of the other eigenvectors of *G*_ss_ to the matrix inverse provide the matrix *G*_qr_ which can be efficiently computed by a rapid convergent matrix series (see [14,15] for details). This point is crucial since it allows for a highly efficient numerical evaluation of all three components of *G*_R_ also for the case where a direct numerical computation of the matrix inverse of (**1** – *G*_ss_) is not possible due to very large values of *N* (note *G*_ss_ has the size *N*_s_ × *N*_s_ with *N*_s_ ≈ *N* ≫ *N*_r_). While *G*_pr_, being typically numerical dominant, has a very simple rank 1 structure the matrix *G*_qr_ contains the most nontrivial information related to indirect hidden transitions. Actually, mathematically both components *G*_pr_ and *G*_qr_ arise from indirect pathways through the scattering nodes (represented by the matrix inverse term in (4)) but *G*_pr_ can be viewed as a uniform background generated by the long-time limit (i.e. the leading eigenvector of *G*_ss_) of the effective process in the complementary scattering network. The component *G*_qr_ gives the deviations from this background and in the following when we speak of “contributions from indirect pathways” we refer essentially to the contributions of *G*_qr_. It is possible that certain matrix elements of *G*_qr_ are negative and if this happens this is also an important information since it indicates a reduction from the uniform background for certain links (matrix elements of *G*_R_, *G*_rr_ and *G*_pr_ are always positive due to mathematical reasons).

Furthermore, we also define the matrix 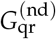 which is obtained from the matrix *G*_qr_ by putting its diagonal elements to zero (these elements correspond to indirect self-interactions of nodes). We consider that this matrix contains the most interesting link information, direct links and “relevant” indirect links describing the deviations from the uniform background due to *G*_pr_. The contribution of each component is characterized by their weights 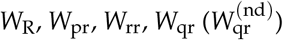 respectively for 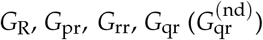. The weight of a matrix is given by the sum of all the matrix elements divided by its size *N*_r_ (*W*_R_ = 1 due to the column sum normalization of *G*_R_). Examples of interesting applications and studies of reduced Google matrices associated to various directed networks are described in [16–19].

### 2.4. Bi-functional Ising MetaCore network

To take into account the bi-functional nature (activation and inhibition) of MetaCore links, we use the approach proposed in [17] with the construction of a larger network where each node is split into two new nodes with labels (+) and (−). These two nodes can be viewed as two Ising-spin components associated to the activation and the inhibition of the corresponding protein. In the construction of the doubled “Ising” network of proteins, each elements of the initial adjacency matrix is replaced by one of the following 2 × 2 matrices

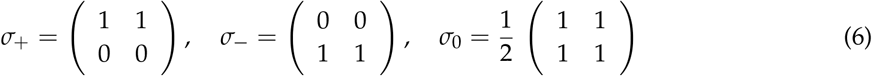

where *σ_+_* applies to “activation” links, *σ_–_* to “inhibition” links, and *σ*_0_ when the nature of the interaction is “unknown” or “neutral”. For the rare cases of multiple interactions between two proteins, we use the sum of the corresponding *σ*-matrices which increases the weight of the adjacency matrix elements. Once the “Ising” adjacency matrix is obtained, the corresponding Google matrix is constructed in the usual way as described above. The doubled Ising MetaCore network corresponds to *N_I_* = 80158 nodes and *N*_*I*,ℓ_ = 939808 links given by the non-zero entries of the used *σ*-matrices.

Now, the PageRank vector associated to this doubled Ising network has two components *P*_+_(*j*) and *P*_–_(*j*) for every node *j* of the simple network. Due to the particular structure of the *σ*-matrices (6), one can show analytically the exact identity, *P*(*j*) = *P*_+_ (*j*) + *P*_–_ (*j*), where *P*(*j*) is the PageRank of the initial single PPI network [17]. The numerical verification shows that the identity *P*(*j*) = *P*_+_ (*j*) + *P*_–_ (*j*) holds up to the numerical precision ~ 10^−13^.

As in [17], we characterize each node by its PageRank “magnetization” given by

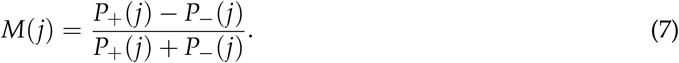

By definition, we have −1 ≤ *M*(*j*) ≤ 1. Nodes with positive *M* are mainly activated nodes and those with negative *M* are mainly inhibited nodes.

In this work the results are mainly presented for the simple network without taking into account the bi-functional nature of links. However, for an illustration we also present some results for the bi-functional network, keeping for further studies a more detailed analysis of this case.

### 2.5. Sensitivity derivative

The reduced Google matrix *G*_R_ of the fibrosis network describes effective interactions between *N*_r_ nodes taking into account all direct and indirect pathways via the global MetaCore network.

As in [9], we determine the sensitivity of PageRank probabilities with respect to a small variation of the matrix elements of *G*_R_. The PageRank sensitivity of the node *j* with respect to a small variation of the link *b* → *a* is defined as

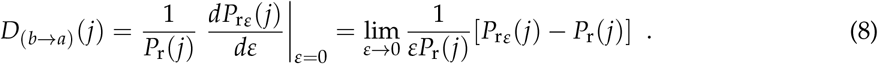

Here, for fixed values of *a* and *b*, *P*_rε_(*j*) is the PageRank vector computed from a perturbed matrix *G*_R*ε*_ where the elements are defined by 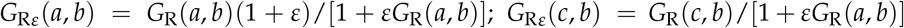 if *c* ≠ *a* and *G*_R*ε*_(*c*, *d*) = *G*_R_(*c*, *d*) if *d* ≠ *b* and for arbitrary *c* (including *c* = *a*). In other words the element *G*_R_(*a, b*), corresponding to the transition *b* → *a*, is enhanced/multiplied with (1 + *ε*) and then the column *b* is resum-normlized by multiplying it with the factor 1/[1 + *εG*_R_(*a*, *b*)] and all other columns *d* ≠ *b* are not modified. We use here an efficient algorithm described in [29] to evaluate the derivative in (8) exactly without usage of finite differences (see also the Appendix for some details on this and other related points). In the following, we consider the case where *j* = *a* and we define the “sensitivity matrix” as 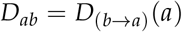. It turns out from the numerical computations that for the cases considered here all values of *D_ab_* are positive: *D_ab_* > 0 which can also be analytically understood as explained in the Appendix.

### 2.6. Determination of external X-proteins

From the experimental results of [5] we have 44 nodes of our selected subset (see first 44 rows of Table 1). Of course, the interactions between these nodes are very important but it is also important to determine how these 44 fibrosis proteins are influenced by external nodes. To find the most important and influential external nodes we take 5 top up- and 5 down-proteins with *K_u_* = 1, …, 5 and *K_d_* = 1, …, 5 from Table 1. Then we determine all external nodes having direct ingoing 134 links to one of these 5 + 5 fibrosis proteins. There are 122 such proteins. The first 44 proteins of Table 1 together with these 122 external proteins (ordered by their PageRank index) constitute an intermediary group of size 166 for which we compute first the reduced Google matrix by (4) and which we note as 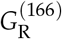 and from this the associated sensitivity matrix 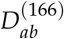 (8) (with *j* = *a*; see also Figure A3). Then we compute the sum of sensitivities 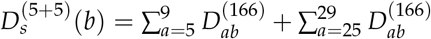 (*a*-sum over top 5 up- and top 5 down-proteins) for *b* = 45, … 166 (new external proteins). Then we select the top 10 external proteins *b* with highest values of 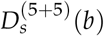. In the following we call this new subgroup the subgroup of X-proteins (or X-nodes). They are given in the last 10 rows of Table 1 (for *K_g_* = 45, …, 54 and *K_x_* = 1, …, 10). We mention that these 10 X-proteins have index values of (1, 2, 3, 4, 6, 8, 10, 15, 27) with respect to the initial list of 122 external proteins (which were already PageRank ordered). It turns out that this procedure selects automatically 10 external nodes which have approximately the strongest PageRank values. This can be understood by the fact that the matrix 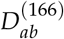 is roughly proportional to *P*(*b*) except for a small number cells with strong peak values (see also Figure A3 and the Appendix for a theoretical explanation). In this way we obtain the full subset of 54 fibrosis proteins given in Table 1. The REGOMAX analysis is performed for these 54 fibrosis proteins and unless stated otherwise all results for *G*_R_, *D_ab_* etc. refer to this group of 54 proteins.

## 3. Results

In this Section we present the results of Google matrix analysis of fibrosis protein-protein interactions.

### 3.1. Fibrosis proteins on PageRank-CheiRank plane

As in [9] we determine the density distribution of all proteins of the MetaCore network on the PageRank-CheiRank plane of logarithms (ln *K*, ln *K*\*) of indexes (*K*, *K*\*) which is shown in Figure 1. The whole plane is divided on 100 × 100 logarithmically equidistant cells and the density is defined as the number of proteins in a given cell divided by a total possible nodes in a given cell (this approach is discussed in more detail e.g. in [24]). The highest density is located at top indexes *K*, *K*\* but in this region there is a relatively small number of proteins. The positions of fibrosis proteins of Table 1 are marked by crosses of 3 colors: red for 10 external X-proteins (*K_x_* = 1, …, 10), pink for 4 TGF-*β* proteins (*K_t_* = 1, 2, 3, 4) and white for the 40 up- and down-proteins (*K_u_*, *K_d_* = 1, …, 20). We see that X-proteins have highest rank positions; 2 of the TGF-*β* proteins approximately follow after *K_x_* values of PageRank and 2 others have significantly lower *K*-rank positions (positions in *K*\*-rank are rather low); proteins *K_u_* and *K_d_* have on average rather low rank positions (very large *K*, *K*\* values). Therefore the X-proteins have the highest network influence and communicativity (small *K*, *K*\* values).

**Figure 1.**
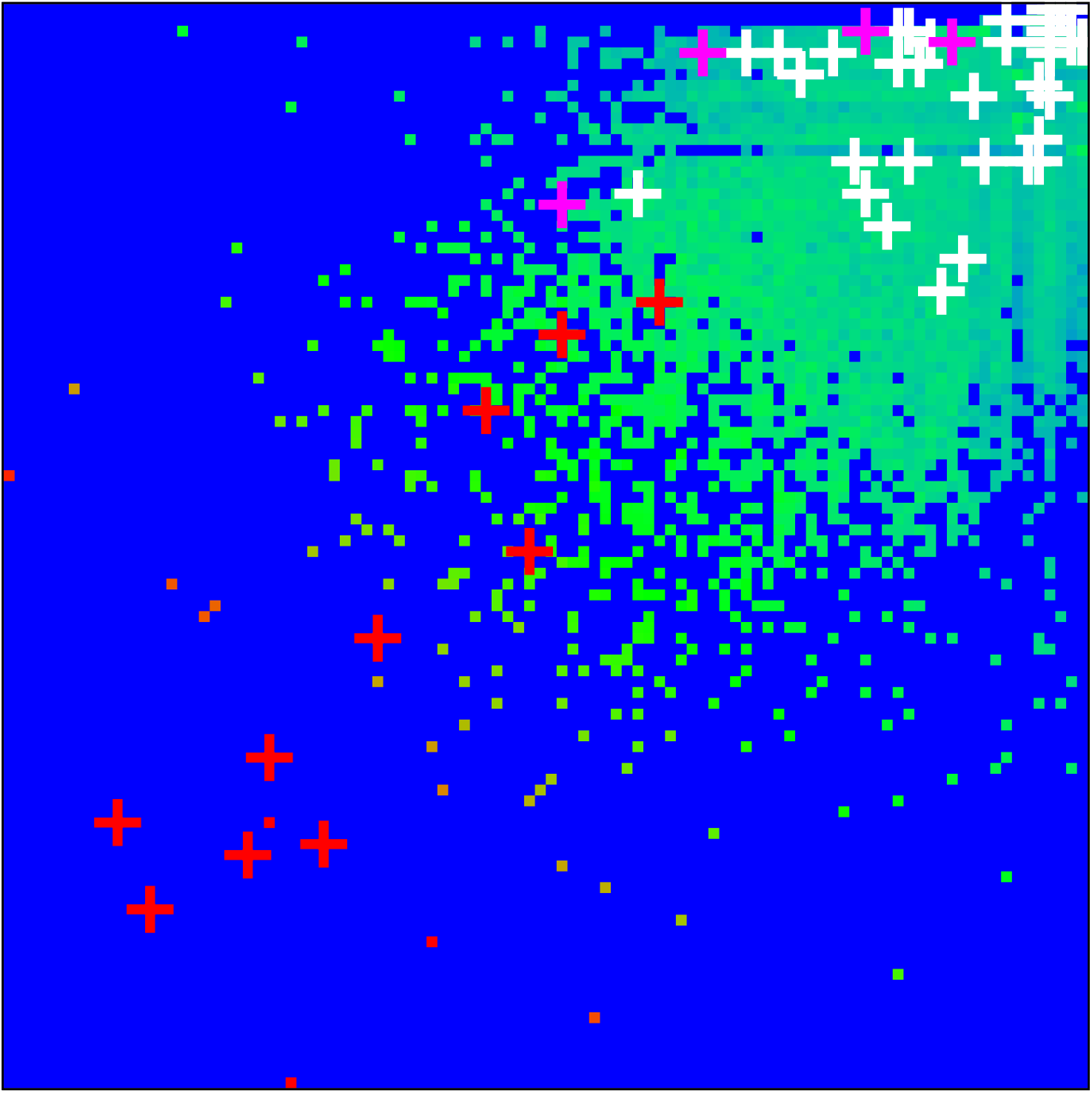
Density of nodes 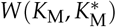 on PageRank-CheiRank plane 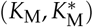 averaged over 100 × 100 logarithmically equidistant grids for 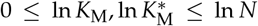, the density is averaged over all nodes inside each cell of the grid, the normalization condition is 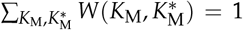. Color varies from blue at zero value to red at maximal density value. In order to increase the visibility large density values have been reduced to (saturated at) 1/16 of the actual maximum density and typical green cells correspond to density values of ~ 1/2^8^ of the (reduced) maximum density. The *x*-axis corresponds to ln *K*_M_ and the *y*-axis to ln 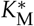 with 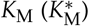 being the global PageRank (CheiRank) index for the full MetaCore network. The crosses mark the positions of the 54 proteins of Table 1 with colors: red for the X-proteins, pink for the TGF-*β* subgroup and white for the up- and down-protein subgroups.

The presentation of Figure 1 uses the global MetaCore rank index values (in the following these values are noted as *K_M_*, *K*\**_M_*; see also Table 1). For the selected subset of 54 fibrosis proteins we note their local rank indexes in this group as *K*, *K*\* which are also given in Table 1. The distribution of these 54 local rank indexes on the PageRank-CheiRank plane of size 54 × 54 is given in Appendix Figure A1.

### 3.2. Reduced Google matrix of fibrosis

The reduced Google matrix *G*_R_ of 54 fibrosis proteins and its 3 matrix components *G*_pr_, *G*_rr_, *G*_qr_ are shown in Figure 2. The weights of these matrices are: *W*_pr_ = 0.9522, *W*_rr_ = 0.0228, *W*_qr_ = 0.0250, 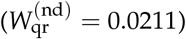 and *W*_R_ = 1 (due to the column sum normalization of *G*_R_). Thus the weight of *G*_pr_ is significantly higher as compared to the two other components. This behavior is quite typical and was also observed for Wikipedia networks (see e.g. [15,18,19]). The physical reason of this is that *G*_pr_ is obtained from the contribution of the leading eigenvector of the matrix *G*_ss_ whose eigenvalue is close to unity and dominates numerically the matrix inverse in (4) (see also the discussion in the last section and [14,15] for details). Furthermore *G*_pr_ has a very simple structure since it is of rank one, i.e. all columns are exact multiples of the first column. Furthermore, these columns are approximately equal to the local PageRank vector. Therefore the component *G*_pr_ does not provide any new interesting information about possible interactions other that it trivially reproduces the PageRank vector.

**Figure 2.**
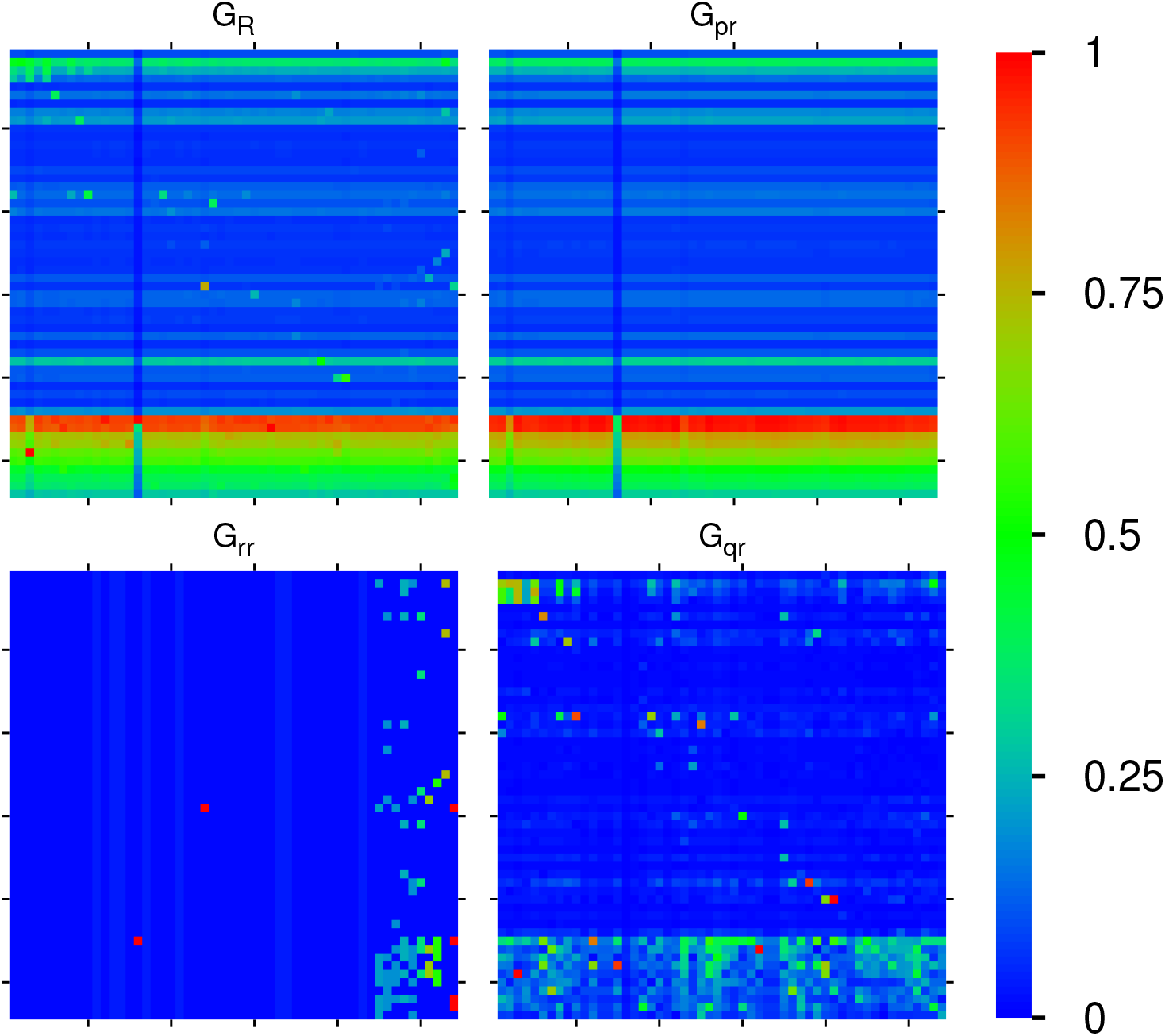
Color density plots of the matrix components *G*_R_, *G*_pr_, *G*_rr_, *G*_qr_ for the group of Table 1; the *x*-axis corresponds to the first (row) index (increasing values of *K_g_*) from top to down) and the *y*-axis corresponds to the second (column) index of the matrix (increasing values of *K_g_* from left to right). The outside tics indicate multiples of 10 of *K_g_*. The numbers in the color bar correspond to 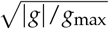 with *g* being the value of the matrix element and *g*_max_ being the maximum value. In order to increase the visibility for the cases of *G*_R_, *G*_rr_, *G*_qr_ the maximum value has been reduced (saturated) to the value of the 3rd largest value of *g* for each case and the cells corresponding to the 1st and 2nd largest values are reduced to the saturation value. In particular *G*_R_(45, 15) (*G*_R_(46, 13)) has been reduced from 0.876387 (0.297512) to *G*_R_(49, 3) = 0.208777; *G*_rr_(45, 16) (*G*_rr_(29, 24)) has been reduced from 0.850004 (0.121432) to *G*_rr_ (29, 54) = 0.019322 (same 3rd value also for the other three cells in column 54); *G*_qr_(49, 3) (G_qr_ (40, 41)) has been reduced from 0.240629 (0.062024) to *G*_qr_ (46, 32) = 0.041108. For the matrix *G*_qr_ there are some negative values and here we show their absolute values (see text).

Numerically *G*_R_ is dominated by *G*_pr_ (with its high weight *W*_pr_ = 0.9521). However, the other two components give us important additional information about direct interactions between the 54 fibrosis proteins (*G*_rr_), and, even more important, about all indirect interactions (*G*_qr_) between these proteins via the global MetaCore network performing an effective summation over all indirect pathways (see [14,15] for details). The weights of the components of *G*_rr_ and *G*_qr_ are comparable. We also see that nearly all direct transitions visible in *G*_rr_ are from X-proteins to other proteins (all subgroups) which is not astonishing due to the selection rule that any X-node must have at least one direct link to the first 5 top- or first 5 up-proteins and also due to the fact the they have rather high PageRank but also CheiRank positions (according to Table 1, Figure 1 and Appendix Figure A1). Since the PageRank probabilities are higher for X-proteins (see Figure 1) there are rather strong transitions to these X-proteins well visible for *G*_R_, *G*_pr_ and to a lesser extent also in *G*_qr_. We note that the component *G*_qr_ has a small number of non-vanishing diagonal matrix elements which appear due to the possibility that a pathway over the global MetaCore network can return to an initial protein.

It should be noted that a few matrix elements of *G*_qr_ have negative values. Such a situation has been already found for other directed networks, e.g. Wikipedia networks studied in [15]. To be more precise for *G*_qr_ and 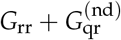 there about 340 out of 2916 negative values (≈ 11%). Most of them are very small. However, there are 10 values between −0.00668 and −0.00334 for both matrices corresponding to 5-10% of the red-color saturation value used for *G*_qr_. However, in Figure 2 only the modulus of matrix elements is shown in order to have a uniform style for all components (the 10 strongest negative values of *G*_qr_ correspond to green color with color bar values of 0.3 to 0.4 and after taking the modulus). Of course, the matrix elements of *G*_R_, *G*_rr_ and *G*_pr_ are always positive due to strict mathematical properties.

Figure 3 shows the effective matrix of transitions for direct links and relevant indirect pathways (without self interactions) which is obtained as the sum of the two components 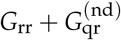. There are also some cells with cyan color for negative matrix elements (corresponding to −0.3 to −0.2 in units of the color bar for the strongest 10 negative values). Most links are due to the interactions from *K_x_* to *K_t_*, *K_u_*, *K_d_* proteins but there are also some other significant transitions between the other members of the group of 54 proteins.

**Figure 3.**
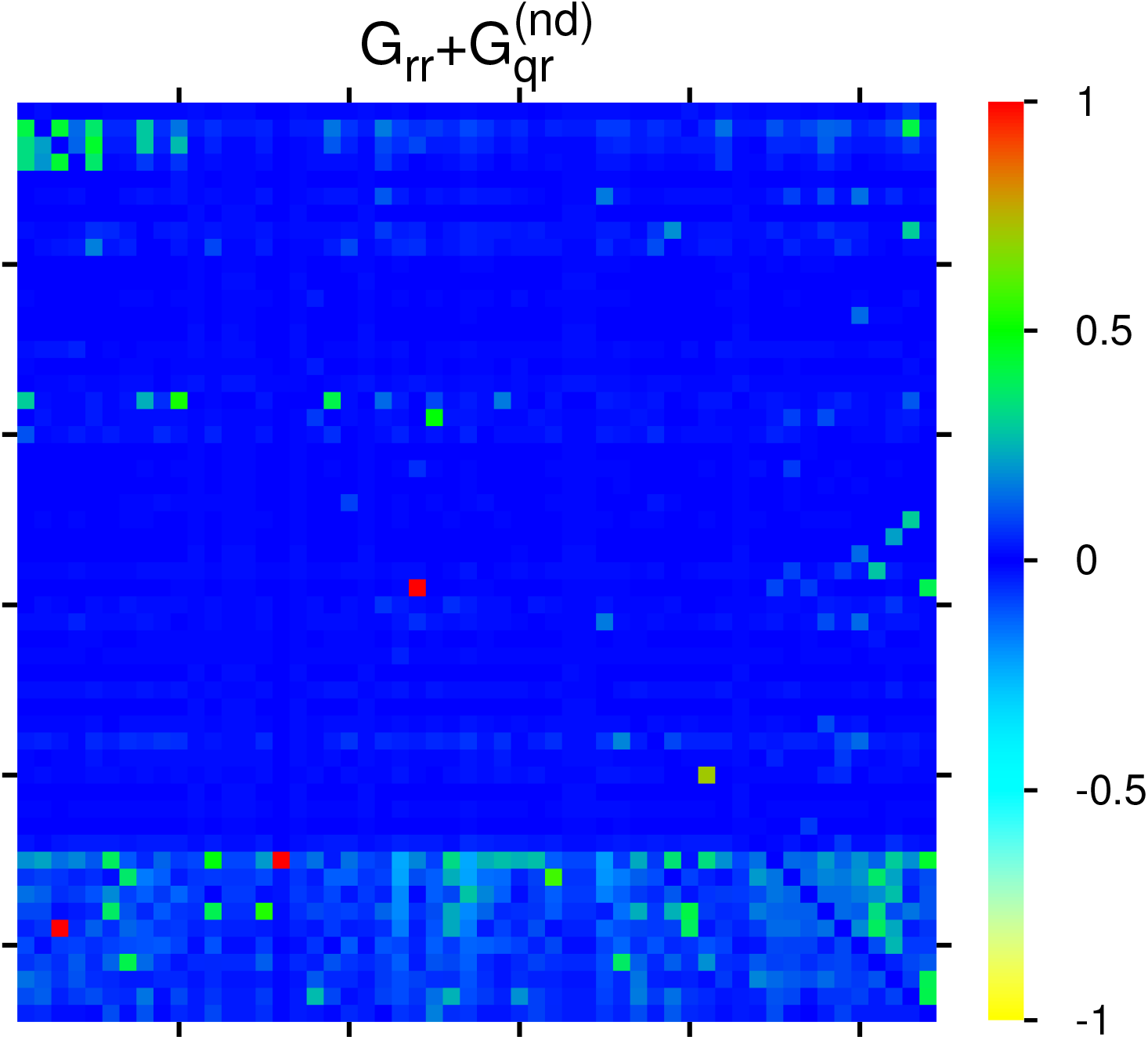
Color density plot 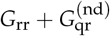 for the group of Table 1 (same plot style and color bar as Figure 2). The matrix element at (45, 16) ((49, 3)) has been reduced from 0.849861 (0.240632) to the value 0.121433 at (29, 24); a few matrix elements of 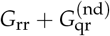 have negative values visible as cyan color (see text).

### 3.3. Network diagrams of fibrosis interactions

In this section we discuss two types of effective networks (of most important PPI links) obtained from the two matrices *G*_R_ and 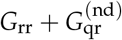, the latter containing the “interesting” links without the uniform background generated by the component *G*_pr_ (and without self-interactions). We remind that the value of a matrix element *g*(*a*, *b*) (with *g* being either *G*_R_ or 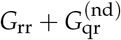) corresponds to the strength of the link *b* → *a*. If this value is sufficiently high we say that *α* is a “friend” of *b* and *b* is a “follower” of *α*. This distinction allows to construct for each matrix two types of effective networks by choosing a few number of “top nodes” and adding a certain number of the strongest friends (or followers) according to the values of |*g*(*a, b*) | and repeating this procedure for a modest number of depth levels.

In Figure 4, we show four graphical representations of such effective networks for the two cases of friend or follower networks and the two matrices *G*_R_ and 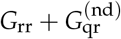 visible in Figures 2 and 3. In these figures and the remainder of this subsection we use the short notations *Tj, Uj, Dj* or *Xj* for a protein/node where *j* = 1, 2, … is the integer value of the subgroup index *K_t_*, *K_u_*, *K_d_* or *K_x_* respectively with real protein names given in Table 1.

**Figure 4.**
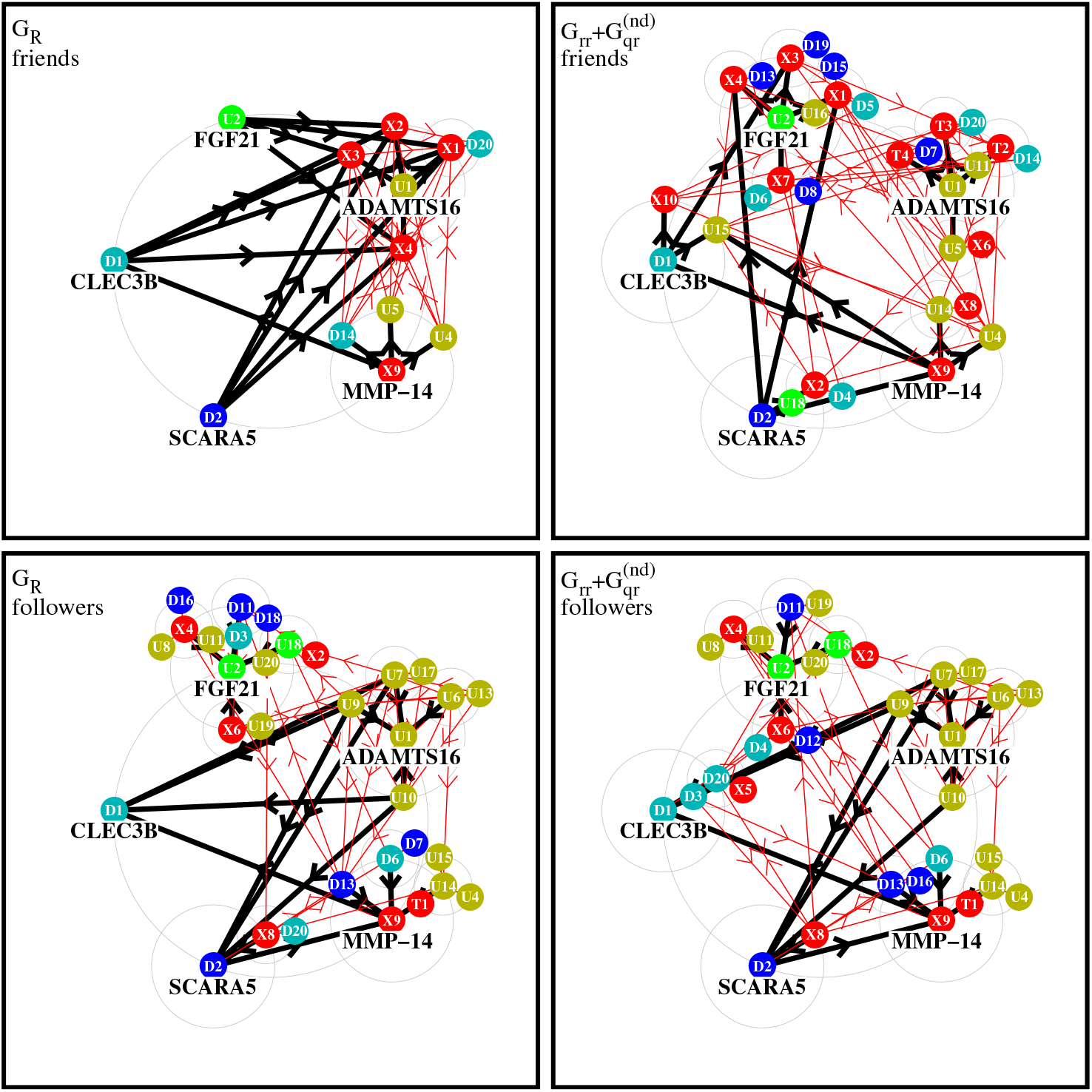
Effective friend and follower networks generated from *G*_R_ and 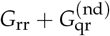. Starting from 5 top nodes the 4 strongest friends/followers for each initial node are selected and links are shown by thick black arrows. For each selected new node further 4 strongest friends/followers are selected and corresponding new links are shown by thin red arrows. In this procedure the direct links between two nodes belonging both to one of the two subgroups of X-proteins or TGF-*β* proteins are not taken into account. The node labels *Tj*, *Uj*, *Dj*, *Xj* (with *j* being an integer value) correspond the local subgroup index *K_t_* = *j*, *K_u_* = *j*, *K_d_* = *j* or *K_x_* = *j* respectively which are given in Table 1. Further details about precise selection rules of links, top nodes and color attribution are given in the text.

To construct the effective network for a matrix component *g* (with *g* being either *G*_R_ or 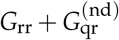) we first choose 5 initial top nodes/proteins corresponding to *U*1, *U*2 (ADAMTS16, FGF21), *D*1, *D*2 (CLEC3B, SCARA5) and X9 (MMP-14). *U*1, *U*2 (*D*1, *D*2) have the strongest positive (negative) TGF-*β* response observed experimentally in [5]. The node corresponding to *X*9 (MMP-14) produces the strongest sensitivity *D_ab_* (among those elements *D_ab_* where *a* is an up- or down protein and *b* is a TGF-*β* or X-protein; see next subsection for details on this). These 5 proteins form the set of level-0 nodes which are placed on a large circle.

We attribute the color red to the combined subgroups of 10 external X-proteins (*K_x_* = 1, …, 10) and 4 TGF-*β* proteins (*K_t_* = 1, 2, 3, 4) The transitions inside this red group are not taken into account since we are mainly interested in the influence of this group on the other up- and down-proteins. We attribute two colors to the up-proteins (olive green to *U*1, green to *U*2) and two colors to the down-proteins (cyan to *D*1, blue to *D*2). Inside the group of up-proteins, we attribute the color olive green to a protein *Uj* if *Uj* is a stronger follower of *U*1 than of *U*2 with respect to 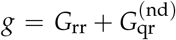, i.e. if *g*(*K_u_* = 1, *K_u_* = *j*) > *g*(*K_u_* = 2, *K_u_* = *j*) and green otherwise. In other words, we compare the strength of the links *Uj* → *U*1 and *Uj* → *U*2 to determine if *Uj* has the color olive green of *U*1 or green of *U*2. In a similar way, comparing the strength of the two links from a *Dj* protein to either *D*1 or *D*2, we attribute the two colors cyan and blue to down-proteins. This attribution rule, using the strongest followers with respect to 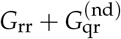 of the two top nodes inside a subgroup, ensures that for all colors there is a considerable number of proteins and it is the same for all four network diagrams (both matrices and both friend/follower cases).

For each of the 5 level-0 proteins, noted *a*, we first search the 4 strongest friends (followers), noted *b*, with largest value of |*g*(*b*, *a*)| (or |*g*(*a*, *b*)|) corresponding the strongest link *a* → *b* (or *b* → *a*), where the matrix *g* is either *G*_R_ or 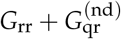. The new nodes *b* (if not yet present in the set of level-0 nodes) nodes form the set of new level-1 nodes and they are placed on medium size circles of level 1 around the corresponding “parent” node *a* of level-0. The links between the nodes *a* and *b* are drawn as thick black arrows with direction *a* → *b* (*b* → *a*) for the friend (follower) case. If a node *b* already belongs to the set of level-0 nodes we also draw a thick black arrow but using its already existing position on the initial large circle. If a node *b* has several parent nodes *a* we place it only on one medium circle preferably around a parent node of the same color if possible.

This procedure is repeated once: for each level-1 protein we determine the 4 strongest level-2 friends (or followers) which are placed on smaller circles of level 2 around the corresponding level-1 protein provided that they are not yet present in the former sets of level-0 or level-1 proteins. The links corresponding to this stage are drawn as thin red arrows with the same directions as in the first stage (we also draw thin arrows for a selected nodes who were already previously selected and using their former positions). As already mentioned above, links where *both* proteins (*a* and *b*) belong to the combined set of X- and TGF-*β* proteins are not taken into account (otherwise they would strongly dominate these diagrams). We limit our-self to two stages of the procedure (i.e three levels of nodes) because otherwise the diagrams would require still smaller circles and many nodes would be hidden by former nodes. We note for the friend-*G*_R_ diagram a further third stage would not add any new nodes since the strongest friends of level-2 are already in the network. For the other cases additional further stages would only add a few number of new nodes with a quite rapid saturation of the network at some limit level where no new nodes are selected.

Figure 4 shows diagrams of level-2 networks for the cases of friend (top row) and follower (bottom row) diagrams and the two matrices *g* = *G*_R_ (left column) or 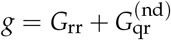 (right column). Concerning the two cases of 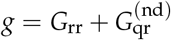 about 15% of the shown arrows correspond to negative values of the matrix element of *g* (link strength is determined by the modulus of the matrix element).

For the friend network of *G*_R_ there is a dominance of links (black arrows) *U*1, *U*2, *D*1, *D*2 → *Xj* for certain X-proteins *Xj* which can be understood by the fact most *Xj* proteins have a significantly higher PageRank probabilities than the other proteins. Furthermore, the total number of nodes in this diagram is quite small because the strongest friends of level-1 nodes (*X*1, *X*2, *X*3, *U*4, *U*5, *D*14) are mostly other level-1 nodes and there is only one new level-2 node (*D*20). This diagram is obviously dominated by the uniform background (of the component *G*_pr_ contributing to *G*_R_) which tends to select mostly the “same new friends” at each level.

For the friend case of 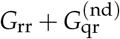 the network structure is significantly richer, since here the global PageRank transitions (due to the uniform background of *G*_pr_) do not play a role. The group around *U*1 includes *T*2, *T*3, *T*4. Thus we see a formation of groups of friends around *U*1 and especially *U*2 with many friends, smaller groups of friends appear around *D*1, *D*2 and *X*9.

For the follower network of 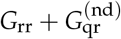, the largest groups of followers are again formed around *U*1, *U*2. In the group around *U*1 we have only other up-proteins while in the group around *U*2 we have up-, down- and X-proteins. The third group around *X*9 is composed of several up- and down-proteins as well as one TGF-*β* protein (*T*1) on level 2. The fourth group around *D*1 includes *D*3, *D*20 and *X*5 but there are also two other followers *U*7, *U*9 but which are placed on the *U*1-circle. The fifth group around *D*2 includes only *X*8 (on its own circle) and *U*7, *U*9, *U*10 from the *U*1-circle.

The follower network of *G*_R_ matrix has a similar structure, since for followers the contribution of *G*_pr_ is not so significant such that several links of followers of *G*_R_ and 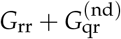 are similar.

It should be noted that the few negative matrix elements of *G*_qr_ have a modest impact on the network diagrams of 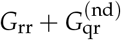 (~ 15% of links and only one stage 1 link for the friend case).

These network diagrams allow to obtain a qualitative graphical view on the most significant fibrosis PPI interactions from a friend or a follower point of view.

We note that in principle it is possible to choose another initial set of 5 proteins at level-0. In Appendix Figure A2, we show the network diagrams for the modified level-0 set: *D*1, *D*2, *U*9, *U*18 and *X*9. Here the 4 up- and down-proteins have the highest sensitivity with respect to X-proteins (see next section). Some features are quite similar as to the first case: the friend diagram of *G*_R_ has only a modest number of nodes with a domination of X-proteins and generally the groups associated to the two up-top nodes appear somewhat larger than the groups for the two down-top nodes.

### 3.4. Sensitivity of fibrosis proteins

In addition to the matrix components *G*_R_, *G*_pr_, *G*_rr_, *G*_qr_ and the network diagrams (of *G*_R_ and 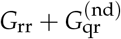 it is also important to analyze the sensitivity matrix *D_ab_* defined previously in (7). This matrix *D_ab_* gives the sensitivity of a protein *a* with respect to a small variation of the transition matrix element of *G*_R_ from protein *b* to *a* on the basis of logarithmic derivative of the PageRank probability (see subsection 2.5 and also the Appendix for more technical details on this).

As described previously (see subsection 2.6), we first compute the sensitivity matrix 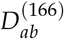 associated to 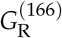 being the reduced Google matrix for a larger intermediary subset containing the 44 TGF-*β*, up- and down-proteins and further 122 external proteins having direct links (of the full MetaCore network) to the first 5 up- (*K_u_* = 1, …, 5) and the first 5 down-proteins (*K_d_* = 1, …, 5). This matrix is shown in Appendix Figure A3.

Then from the set of 122 external proteins we select the 10 proteins *b* with the largest effective sensitivity given by the sum 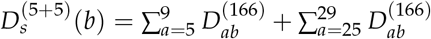 (see Subsection 2.6) which form the group of 10 X-proteins. The 44 TGF-*β*, up- and down-proteins together with these 10 X-proteins form our main group of 54 proteins given Table 1 and for which have presented results of the reduced Google matrix in the last subsections.

The sensitivity matrix *D_ab_* of size 54 × 54 for this main group is shown in Figure 5 with zoomed parts visible in Figure 6.

**Figure 5.**
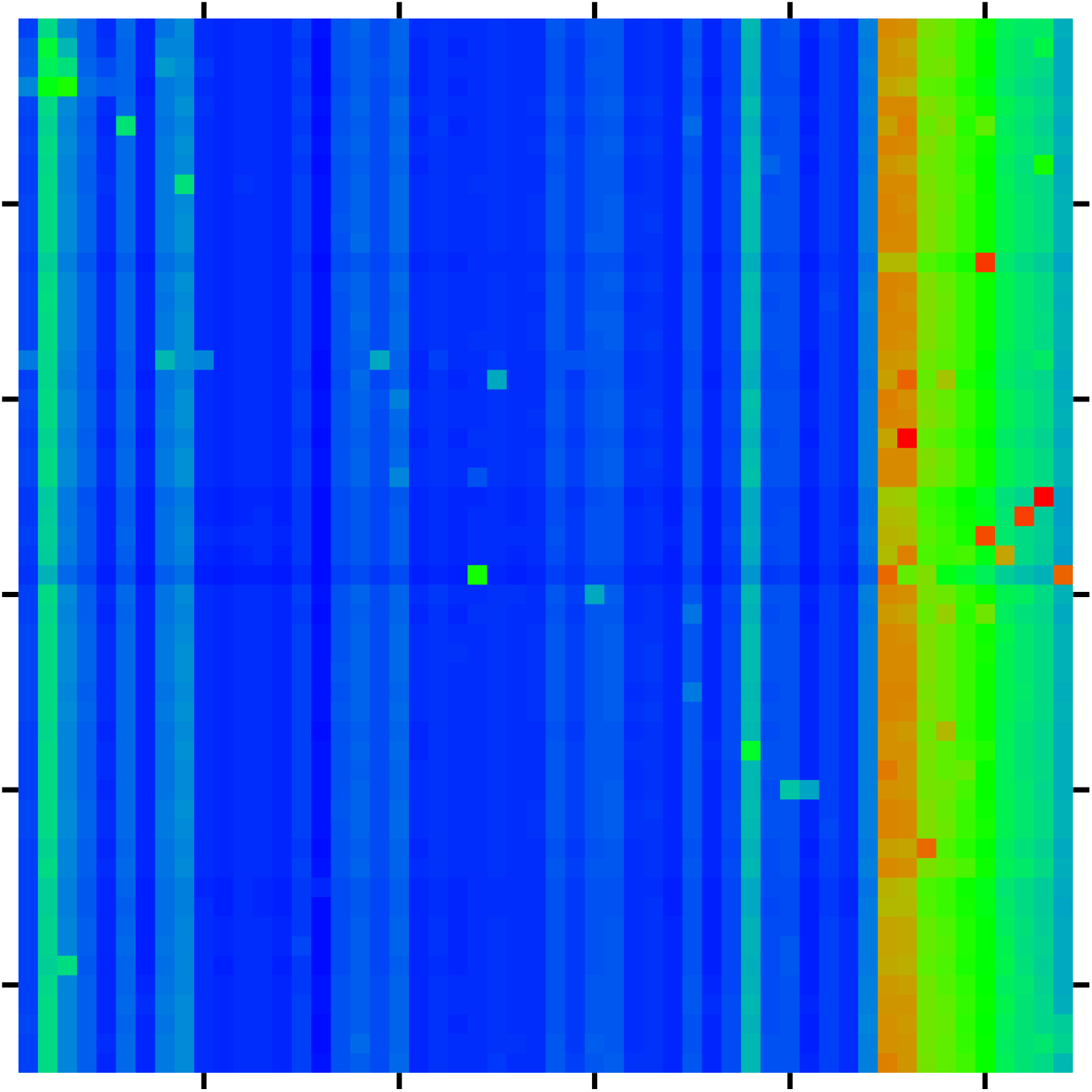
Color density plot of the sensitivity matrix *D_ab_* of fibrosis proteins of Table 1; the axes and colors are defined as in Figure 2 (without saturation); the strongest top 40 sensitivity values are given in Table 2.

**Figure 6.**
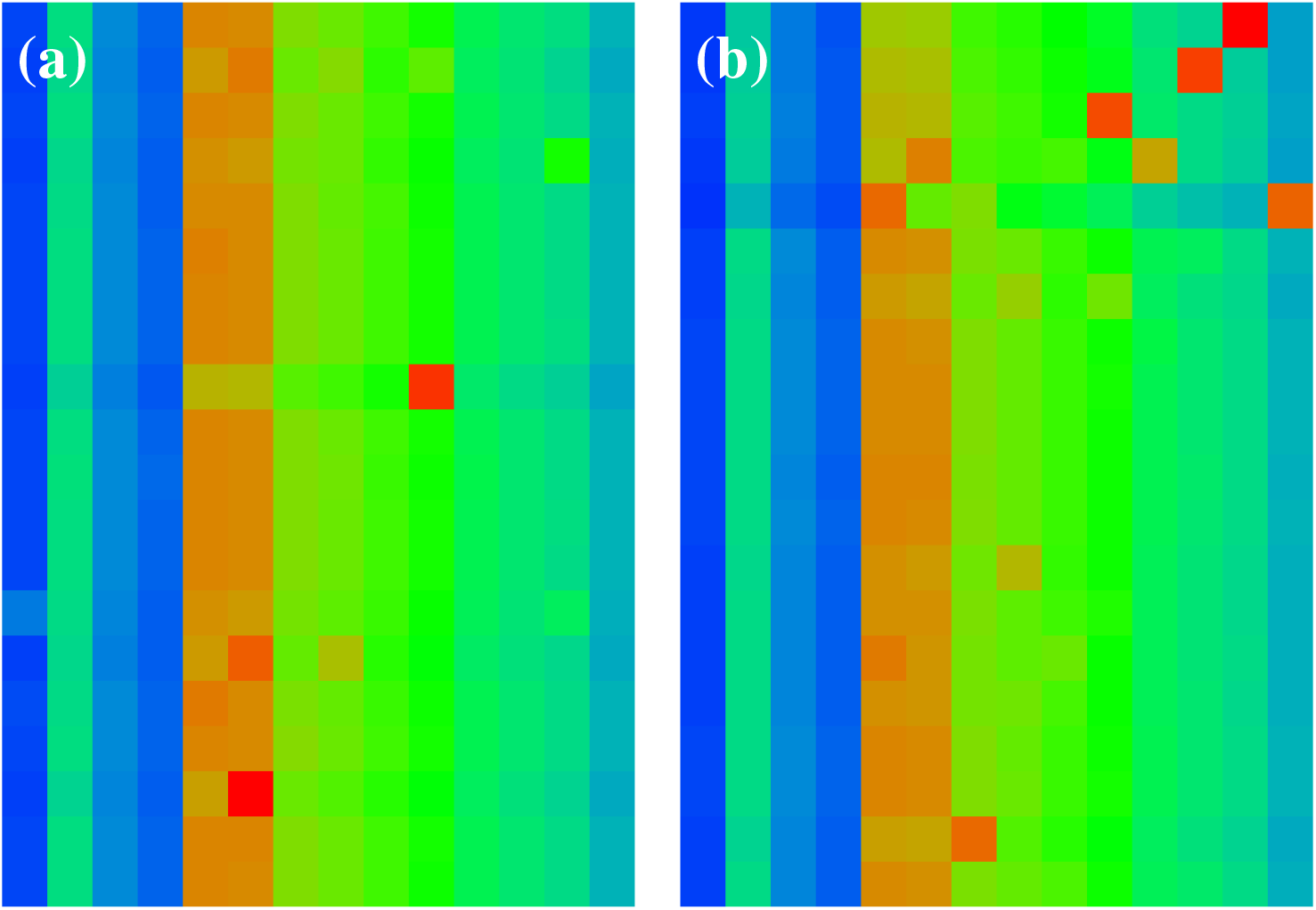
Zoomed parts of sensitivity matrix *D_ab_* of Figure 5. Both panels show a selected sub-region of Figure 5 with the index *a* (vertical axis from top to down) belonging to the set of up-nodes (*a* = 5, …, 24 in panel (a)) or down-nodes (*a* = 25, …, 44 in panel (b)) and the index *b* (horizontal axis from left to right) corresponds for both panels to the 4 nodes of the TGF-*β* subgroup (*b* = *K_t_* = 1, … 4 for 4 left columns in each panel) and the 10 nodes of the *X*-proteins (*b* = 45, … 54 or *K_x_* = 1, …, 10 for 10 right columns in each panel).

We mention that the appearance of MMP-14 (*K_x_* = 9) at the top position of Table 2 is the reason why we selected this protein as one of the 5 top nodes in the net diagrams discussed in the last subsection. For the net diagrams shown in Figure 4, the other four top nodes were simply chosen as the first two up- (*K_u_* = 1, 2) and down-proteins (*K_d_* = 1, 2). However, for the net diagrams shown in Appendix Figure A2 also the two top up- and down-nodes were chosen by the criterion of top positions in Table 2 resulting in *K_u_* = 9, 18 and *K_d_* = 1, 2.

**Table 2.**
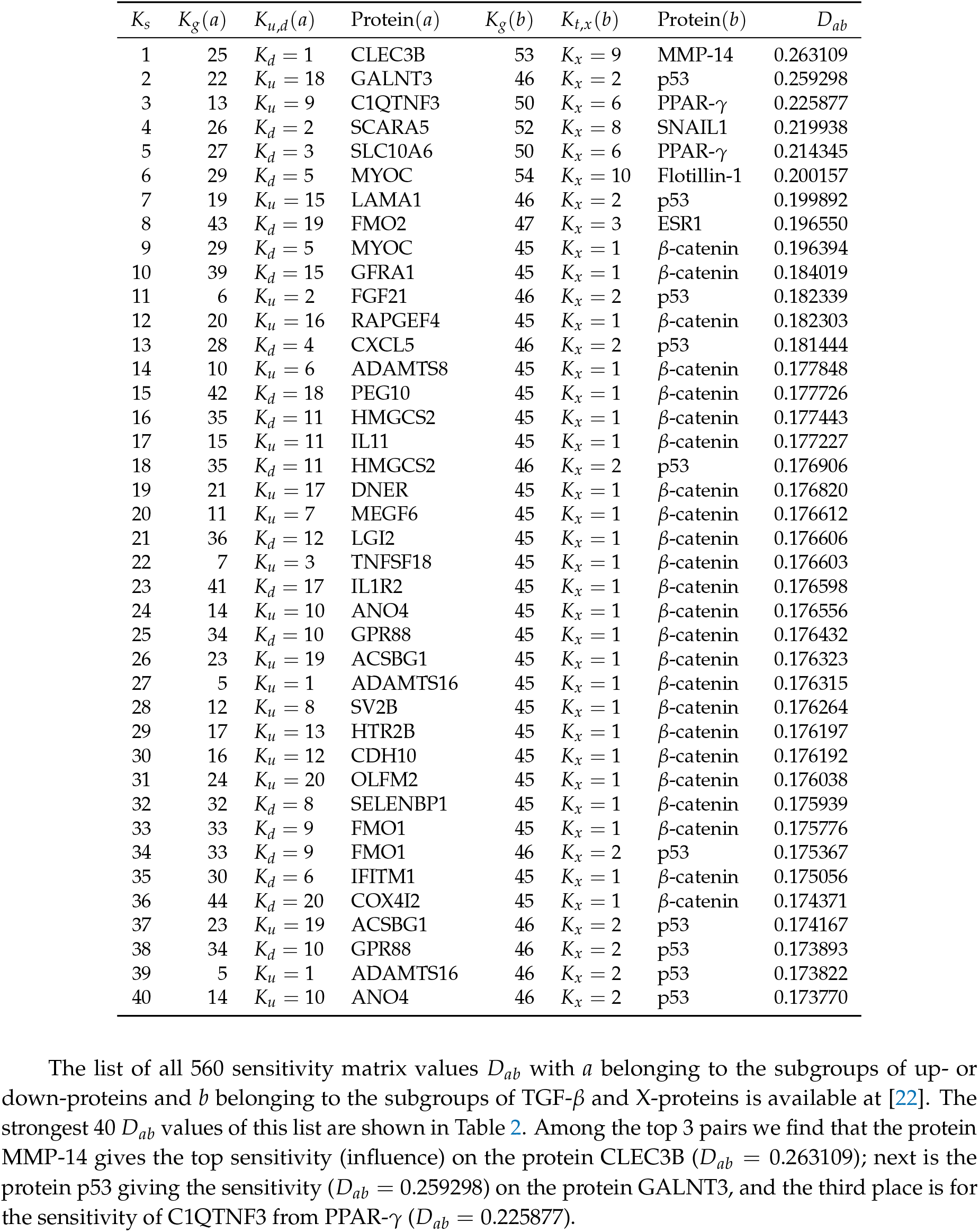
List of 40 top protein pairs (*a*, *b*) with strongest sensitivity matrix element *D_ab_* with *a* belonging to the subgroups of up- or down-proteins and *b* belonging to the subgroups of TGF-*β* and X-proteins. The first column gives the ranking index *K_s_* of *D_ab_* matrix elements ordered by a decreasing value, the 2nd to 4th columns provide the *K_g_*, *K_u,d_* indexes and the name of the protein (*a*), the 5th to 7th columns provide the *K_g_*, *K_t,x_* indexes and the name of the protein (*b*), and the 8th column shows the value of *D_ab_*. See also Figure 5 which shows a color density plot for all matrix elements *D_ab_* and Table 1 for the list of considered proteins. An ordered list of all 560 values of sensitivity influence values *D_ab_* of TGF-*β* or X-proteins (for “*b*”) on up-/down proteins (for “*a*”) is available at [22].

We have also computed the effective TGF-*β* sensitivity on up- or down-proteins (noted *a*) defined by the sum 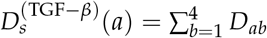. Ordering these values in decreasing order we obtain the ranking index 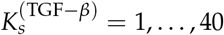 whose dependence on *K_u_* and *K_d_* is visible Appendix Figure A4. We see that for the up-proteins we have 14 ranking values located at 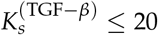 and for the down-proteins only 6 values at 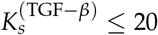 (with 3 values at 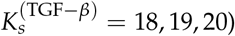. This shows that the overall influence of TGF-*β* proteins is somewhat stronger on the up-proteins as compared to the down-proteins.

However, we mention that the different values of 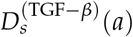 used to determine this ranking have only modest size variations in the interval 0.0250 to 0.0465 with most values between 0.040 and 0.043. Furthermore, in global the external X-proteins have a much higher influence (on up- and down-proteins) than the TGF-*β* proteins. For instance in Table 2 the TFG-*β* proteins do not appear at all (in the three “*b*” columns) and in the full list of 560 entries the first appearance of a TFG-*β* protein is at the ranking position *K_s_* = 319.

Both of these points can be explained by the approximate expression *D_ab_* ≈ [1 – *P*_r_(*a*)]*P*_r_(*b*) ≈ *P*_r_ (*b*) which is derived in the appendix for a simplified model of a rank 1 *G*_R_ matrix but which also holds approximately for arbitrary *G*_R_ matrices due to the strong numerical weight of the rank 1 component *G*_pr_. This behavior is also confirmed, for a “uniform background”, by Figures 5 and 6 for *D_ab_* and Appendix Figure A3 for 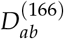. However, there are typically some exceptional peaks at a few values of the (*a, b*) index pair where strong deviations from this simple expression are possible and which are due to the components of *G*_rr_ and *G*_qr_ in *G*_R_.

Essentially *D_ab_* ~ *P*_r_(*b*) does not (strongly) depend on *a* explaining that the values of the partial sum 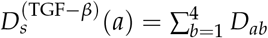 show only modest size variations. Furthermore, Table 2, containing the largest *D_ab_* values (with *b* being either an X or a TGF-*β* protein and *a* being an up- or down protein), is dominated by X-proteins which have mostly larger *P*_r_(*b*) values than the TGF-*β* proteins.

We also determine the global influence on the whole group of fibrosis up- and down-proteins by computing the sum 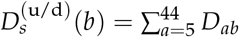 (i.e. the *a*-sum is over up- and down-proteins) for each X or TGF-*β* protein *b*. The resulting values of this quantity are provided in Table 3. According to the simple expression for *D_ab_* we have a linear dependence of 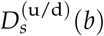 on *P*_r_(*b*) and due to the a-sum the effect of exceptional peaks is strongly reduced. This linear dependence is clearly visible in Table 3 and Appendix Figure A5. A simple linear fit 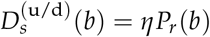 provides the value *η* = 39.5 ± 1.4 for the coefficient and a more general power law fit 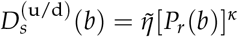 results in a similar coefficient 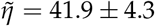 and an exponent *κ* = 1.017 ± 0.028 close to unity.

**Table 3.** Values of the sum 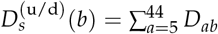 (i.e. the *a*-sum is over up- and down-proteins) for *b* belonging to the TGF-*β* or the X-proteins subgroups. The list is ordered with respect to decreasing 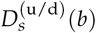 values with the first column giving the corresponding ranking index; the 2nd and 3rd columns giving the *K_g_*, *K_t,x_* indexes; the 4th and 5th columns containing the local PageRank index *K* and the name of the protein *b* and the 6th and 7th columns giving the values of 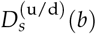 and the local PageRank probability *P*_r_ (*b*). Both *K* and *P*_r_ (*b*) correspond to the group of 54 fibrosis proteins of Table 1.

| Rank | $K_g(b)$ | $K_{t,x}(b)$ | $K$ | Protein($b$) | $D_s^{(u/d)}(b)$ | $P_r(b)$ |
| --- | --- | --- | --- | --- | --- | --- |
| 1 | 45 | $K_x = 1$ | 1 | β-catenin | 6.809993 | 0.175768 |
| 2 | 46 | $K_x = 2$ | 2 | p53 | 6.789229 | 0.171249 |
| 3 | 47 | $K_x = 3$ | 3 | ESR1 | 4.513399 | 0.113285 |
| 4 | 48 | $K_x = 4$ | 4 | STAT3 | 4.109638 | 0.104088 |
| 5 | 49 | $K_x = 5$ | 5 | RelA | 3.343309 | 0.085443 |
| 6 | 50 | $K_x = 6$ | 6 | PPAR-γ | 3.086237 | 0.070668 |
| 7 | 51 | $K_x = 7$ | 7 | IKK-β | 1.696330 | 0.043249 |
| 8 | 52 | $K_x = 8$ | 8 | SNAIL1 | 1.477019 | 0.034269 |
| 9 | 53 | $K_x = 9$ | 10 | MMP-14 | 1.368302 | 0.029121 |
| 10 | 2 | $K_t = 2$ | 9 | TGF-β 1 | 1.081828 | 0.029166 |
| 11 | 54 | $K_x = 10$ | 12 | Flotillin-1 | 0.787569 | 0.016863 |
| 12 | 3 | $K_t = 3$ | 13 | TGF-β 2 | 0.333981 | 0.012451 |
| 13 | 4 | $K_t = 4$ | 20 | TGF-β 3 | 0.159633 | 0.004157 |
| 14 | 1 | $K_t = 1$ | 30 | TGF-β 0 | 0.081261 | 0.002090 |

However, Table 3 also shows that at the ranking positions 9 (*K_x_* = 9 for MMP-14) and 10 (*K_t_* = 2 for TGF-*β* 1) there is one ranking inversion between 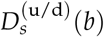 and *P*_r_ (*b*). The value of 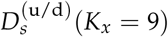 is roughly 30% larger than 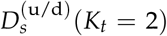 while the PageRank value of the former is very slightly (0.15%) smaller than the value of the latter (both PageRank values are nearly identical). In Appendix Figure A5 both of these proteins correspond to two data points with a certain visible (vertical) difference for 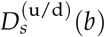 but with no visible (horizontal) difference for *P*_r_(*b*).

We argue that the obtained high sensitivity values shown in Figures 5, 6 and Table 2 can be tested in experiments similar to those reported in [5]. Also the global influence 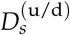 from Table 3 gives us a prediction of the globally stronger influence of the X-proteins than the TGF-*β* proteins. These results open new perspectives for external proteins influence on fibrosis.

### 3.5. Bi-functional of fibrosis network

Here we present in short certain results for the bi-functional MetaCore network. The doubled Ising MetaCore network has *N_I_* = 80158 nodes and *N*_*I*,ℓ_ = 939808 links. We compute the reduced Google matrix *G*_R_ for the doubled number of nodes 2 × 54 = 108 (by attributing (+) and (−) labels to each node) for the fibrosis proteins of Table 1. Here we present only some selected characteristics, all data for the Ising Google matrix are available at [22].

In Figure 7 we show the magnetization *M*(*j*) = (*P_+_*(*j*) – *P*_–_(*j*))/(*P*_+_(*j*) + *P*_–_(*j*)) of proteins of Table 1 with their location on PageRank-CheiRank plain (*K*, *K*\*). We remind that *P*_±_ (*j*) is the PageRank value of the node *j* with label (±) and that the sum satisfies *P*(*j*) = *P*_+_(*j*) + *P*_–_(*j*) where *P*(*j*) is the PageRank value of the node *j* of the simple network. The magnetization is positive for nodes which are more likely to be activated or in other words which have on average more incoming activation links (and/or coming from other nodes with larger PageRank values) than inhibition links while negative values correspond to nodes being more likely to be inhibited by other nodes.

**Figure 7.**
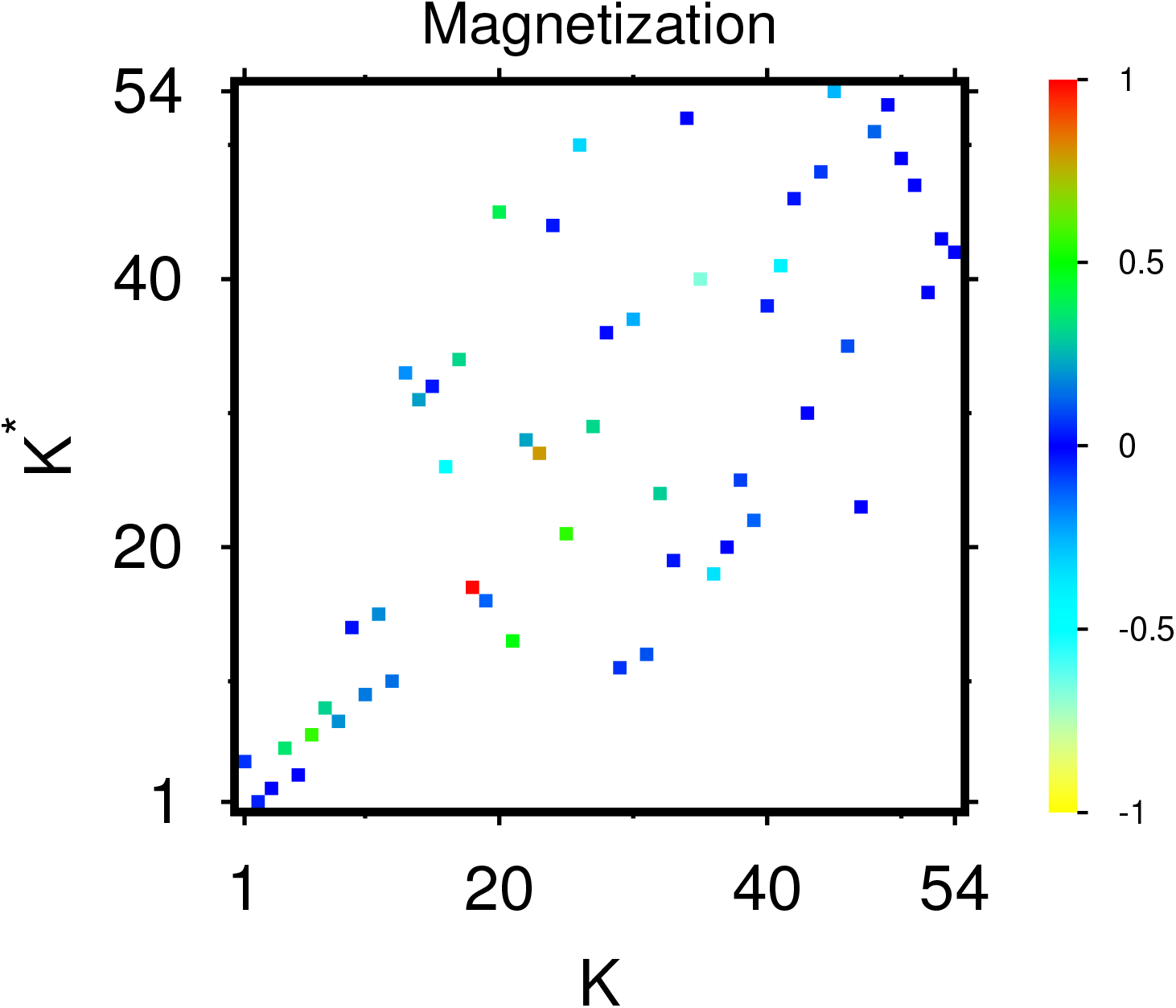
PageRank “magnetization” *M*(*j*) = (*P*_+_ (*j*) – *P*_–_(*j*)) / (*P*_+_ (*j*) + *P_–_*(*j*)) of proteins of Table 1 shown on the PageRank-CheiRank plane (*K*, *K*\*) of local indices; here *j* represents a protein node in the initial single protein network and *P*_±_ (*j*) are the PageRank components of the bi-functional Ising MetaCore network (see text). The values of the color bar correspond to *M/* max |*M*| with max |*M*| = 0.690937 being the maximal value of | *M*(*j*) | for the shown group of proteins. Note that the positions in PageRank-CheiRank plane are identical to the positions of Appendix Figure A1 and the corresponding *K*, *K*\* values are given in the 3rd and 4th column of Table 1.

According to Figure 7 the majority of proteins have values of *M* being close to zero (neutral action on average coming from other nodes) but also there are some nodes with with significant positive values such as RAPGEF4 (at *K* = 18, *K*\* = 17, *K_g_* = 20, *K_u_* = 16, *M* = 0.690937) corresponding to the only red box (maximum value of 1 in units of the color bar) and HMGCS2 (at *K* = 23, *K*\* = 27, *K_g_* = 35, *K_d_* = 11, *M* = 0.550286) with an orange-brown box (value of 0.8 in units of the color bar). There are about further 9 proteins with various degrees of green color (*M* values between 0.2 and 0.4 corresponding to 0.3 to 0.6 in units of the color bar). The two proteins with strongest negative values of *M* are CLEC3B (at *K* = 35, *K*\* = 40, *K_g_* = 25, *K_d_* = 1, *M* = −0.463912) with a light cyan box (value of −0.7 in units of the color bar) and ACAN (at *K* = 16, *K*\* = 26, *K_g_* = 8, *K_u_* = 4, *M* = −0.342585) with a cyan box (value of −0.5 in units of the color bar). There are about 5 further proteins with various degrees of cyan color (M values between −0.28 and −0.17 corresponding to −0.4 to −0.25 in units of the color bar). We note that CELC3B is also selected in both network diagrams of Figure 4 and Appendix Figure A2 as one of the two down-top-nodes either because it is the first protein in the list of down-proteins or because it appears at the top position of Table 2 for the strongest sensitivity value *D_ab_* (with *α* being CELC3B and *b* being the X-protein MMP-14). One may also note that Appendix Figure A1 shows the same (*K*, *K*\*) positions as Figure 7 and allows to identify which of the boxes belong to the subgroups of TGF-*β* proteins, up- or down-proteins, or X-proteins, The complete table of magnetization values used for Figure 7 including the values of *K*, *K*\*, *K_g_* etc. is available in one of the data files provided in [22].

In Figure 8 we show the matrices components *G*_R_ and 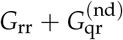 for the group of selected 108 nodes corresponding to the Ising MetaCore network. Their structure is quite similar to the corresponding components for the group of 54 nodes for the simple network shown in Figures 2 and 3, i.e. *G*_R_ is dominated by the uniform background due to the component *G*_pr_ with some exceptional peak values and large values if the first (vertical) matrix index correspond to an X-protein with large PageRank probability. For 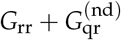 the structure is more sparse showing the most significant direct and relevant indirect transitions. We note that for the Ising case, the matrix values are identical for the two labels of given node in horizontal position (except for the diagonal elements of 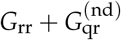 which have been artificially put to zero) which is a mathematical property of these matrices. However, in vertical direction there are significant differences between the two Ising labels especially for 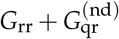.

**Figure 8.**
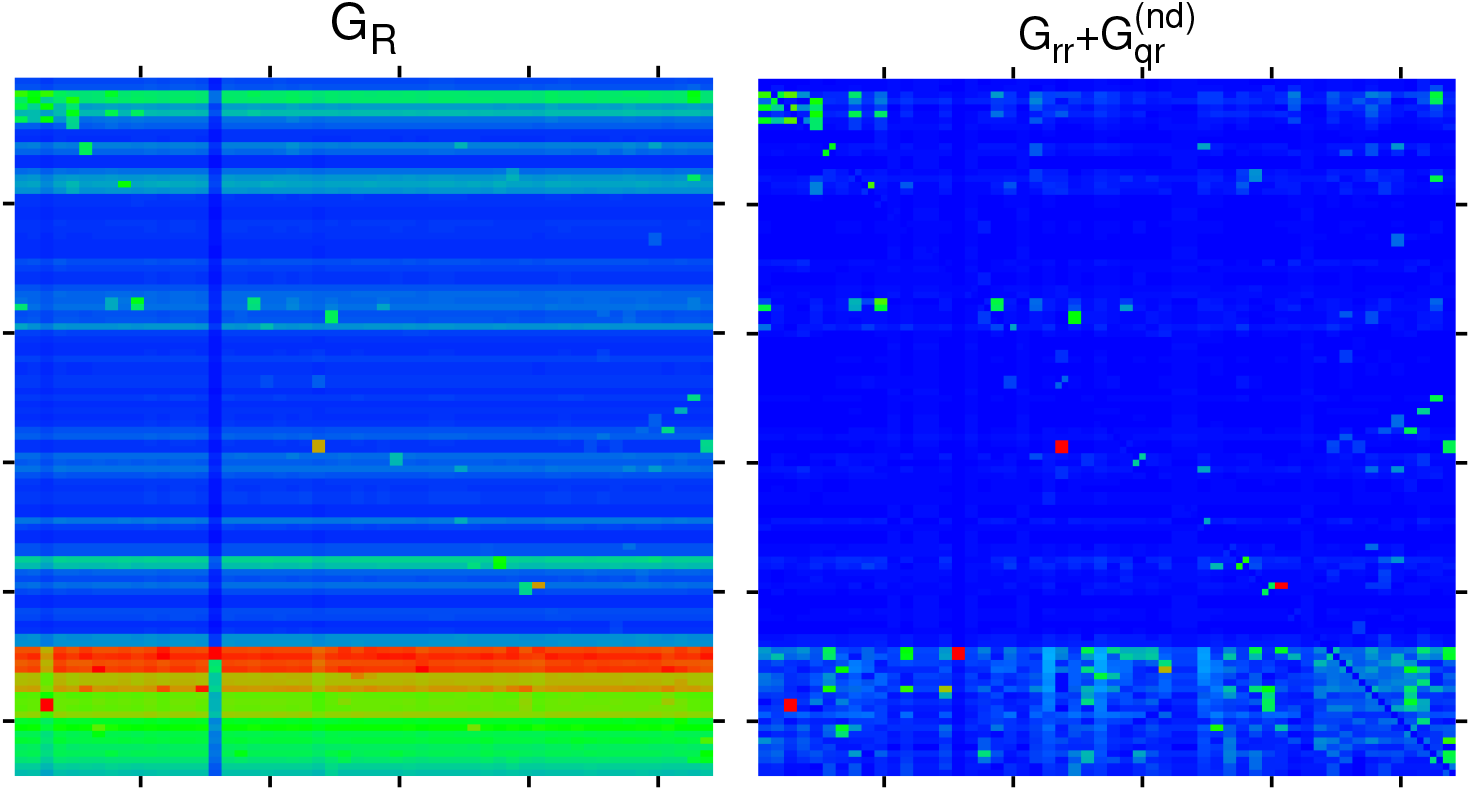
Color density plots of *G*_R_ and 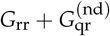 for the bi-functional Ising MetaCore network and the extended group of 108 nodes by attribution of labels (+) and (−) to each node of Table 1. The matrix plot style is similar as in Figure 2 with outside tics indicating multiples of 20 of the index values. The color bar is as in Figure 2 with the same translation of colors to matrix values. The saturation value is for both panels the 6th largest value for each matrix and larger values are reduced to this value. The strongest cell values are reduced from 0.437575 (0.424939) to 0.101874 (0.060717) for 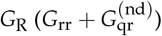.

Further detailed analyses of the Ising MetaCore network with applications on fibrosis interactions are kept for future studies. However, an interested reader can find additional numerical results at [22]. In particular figures for the Ising network diagrams obtained from the Ising versions of *G*_R_ and 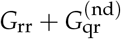 in the same way as in subsection 3.3 are available there.

## 4. Conclusion

In this work we presented the Google matrix analysis of protein-protein interactions of fibrosis. The group of 54 proteins actively participating in the fibrosis process is determined on the basis of INSERM experimental results presented in [5] which give 44 proteins and in addition we add 10 external proteins with strongest sensitivity (on certain 10 proteins of the 44-group) which is computed in the context of the REGOMAX approach applied to the MetaCore network [6]. Our results allow to identify the most important interactions between this fibrosis group of 54 proteins. We hope that these algorithms allow to predict those proteins which will produce a significant influence on the fibrosis process. We expect that future experiment will allow to check these predictions obtained from the Google matrix analysis.

## Appendix

### A1. Additional figures for REGOMAX results

Here we present additional Appendix Figures A1, …, A5 for the main part of this article. Appendix Figure A1 provides complementary information to Figure 1.

**Appendix Figure A1.**
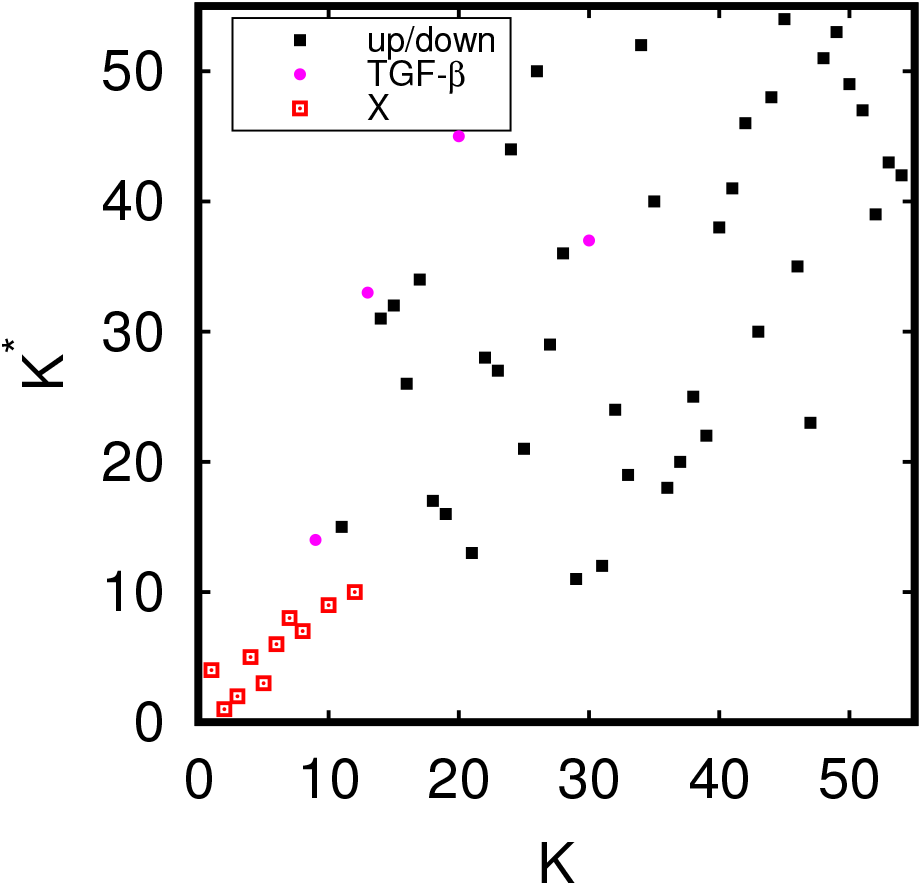
Positions of the 54 proteins of Table 1 in the local PageRank-CheiRank. Note that these positions are identical to the positions of Figure 7 and the corresponding *K*, *K*\* values are given in the 3rd and 4th column of Table 1. Pink full circles correspond to the subgroup of TGF-*β* nodes, full black boxes correspond to the subgroups of up- and down-proteins and red squares correspond to the subgroup of X-proteins.

**Appendix Figure A2.**
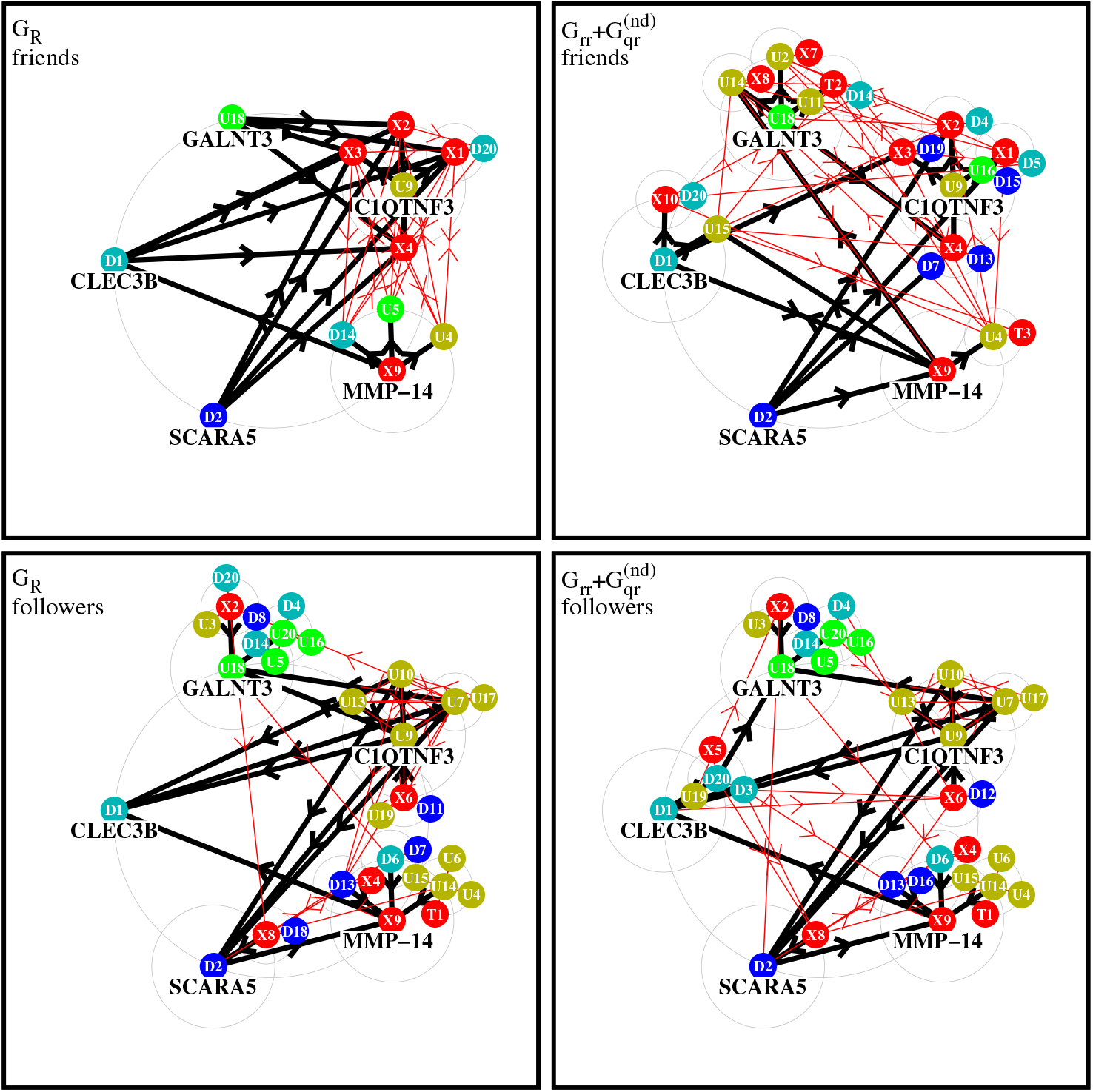
Effective network diagram for the same cases as in Figure 4 but using different 5 top nodes being the first X-node, the first two up-nodes and the first two down-nodes according to Table 2.

Appendix Figure A2 provides the network diagrams similar as in Figure 4 but with a different choice of 5 top nodes based on the criterion of top positions in Table 2.

**Appendix Figure A3.**
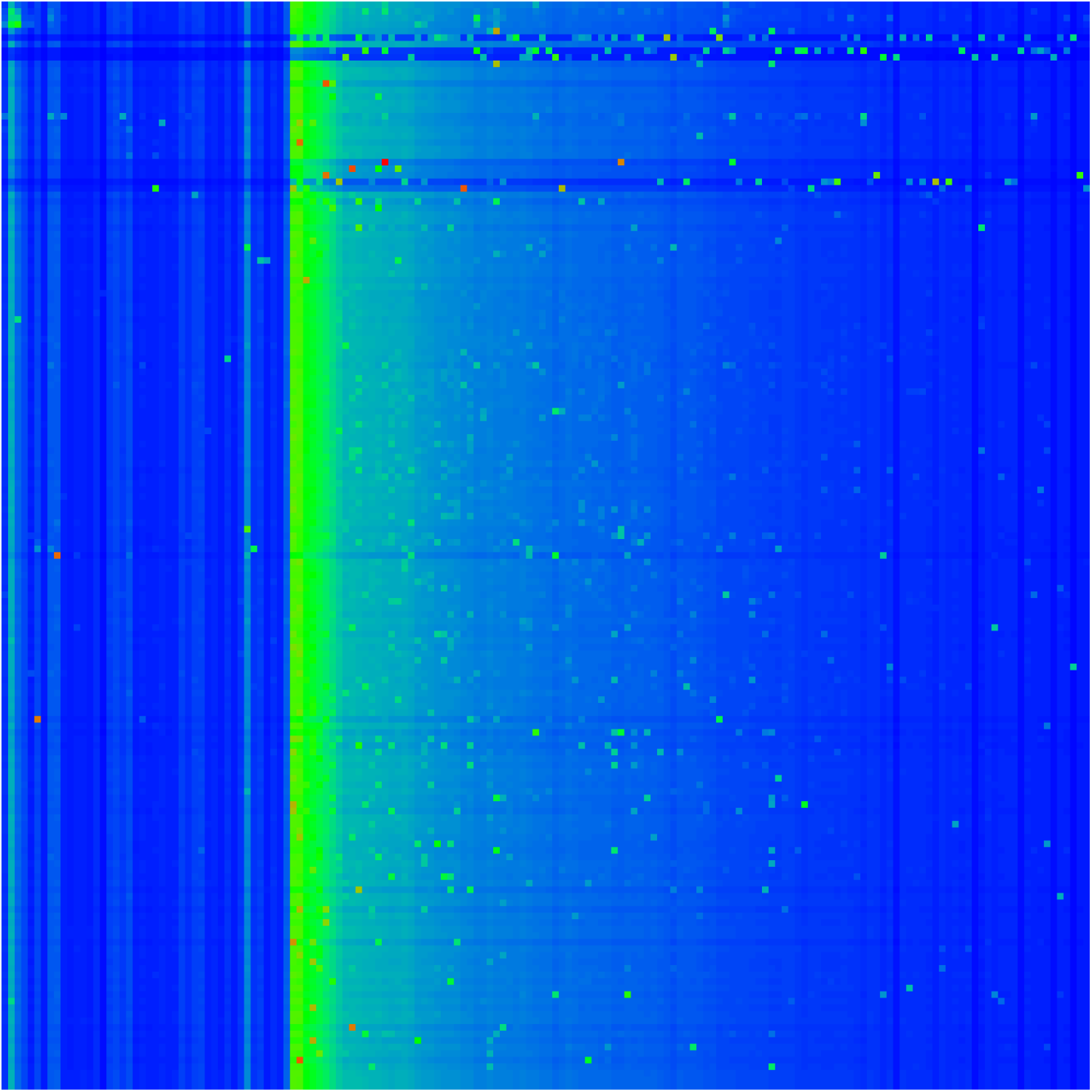
Color density plot of the sensitivity matrix 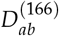 for the intermediary group of 166 proteins being the first 44 proteins of Table 1 (TGF-*β*, up- and down-subgroups) and 122 further proteins (in PageRank order) determined by having a direct link to one of the top 5 up-nodes (*K_u_* ≤ 5) or top 5 down-nodes (*K_d_* ≤ 5; see also text).

Appendix Figure A3 shows the sensitivity matrix 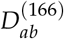 for the intermediary group of 166 proteins which was used to determine the additional 10 X-proteins as explained in subsection 2.6.

**Appendix Figure A4.**
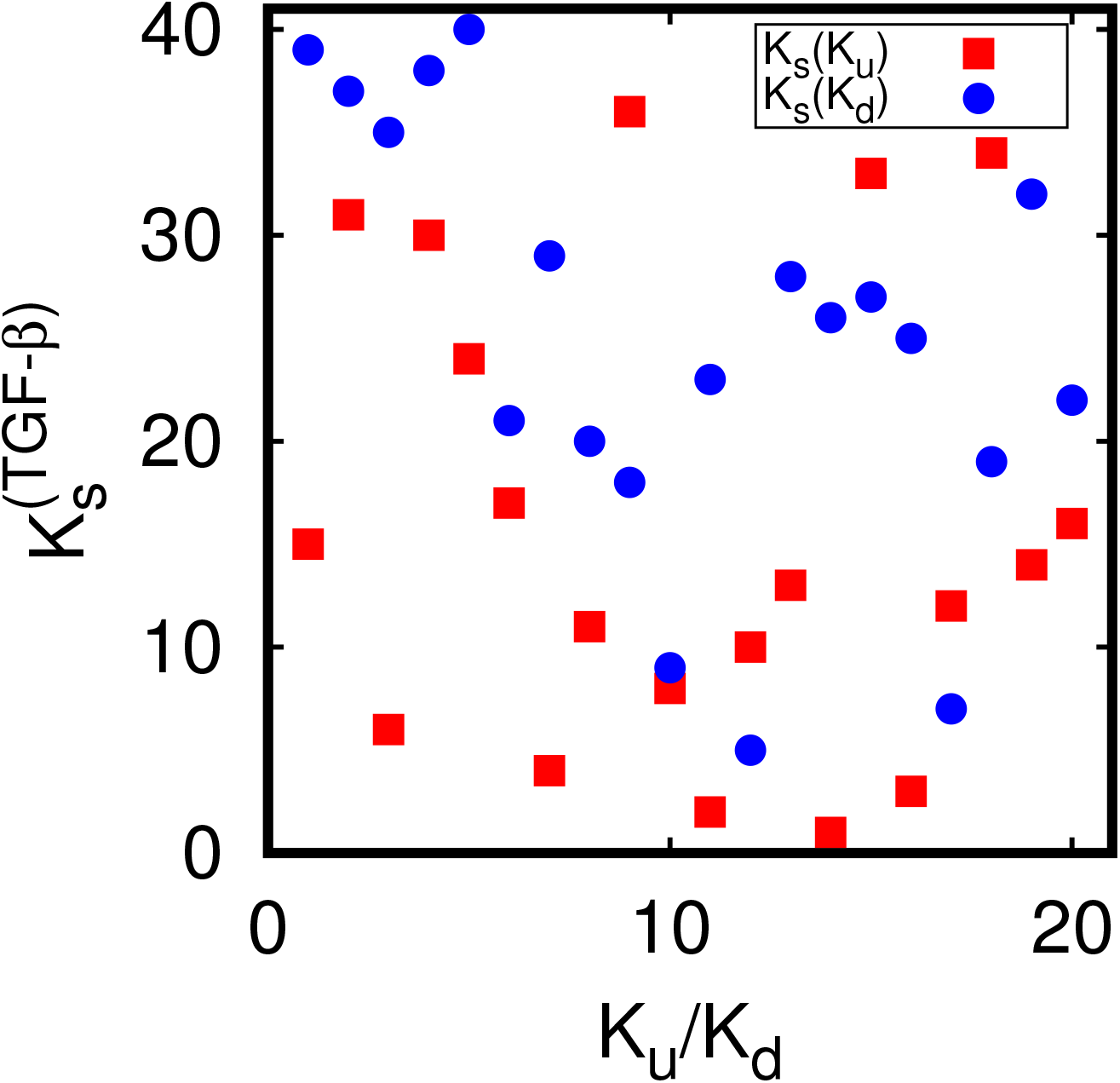
Effective ranking 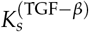 index of the TGF-*β* sensitivity versus *K_u_ /K_d_* of up- (red boxes) and down-proteins (blue full circles). The ranking index 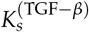 is determined by ordering the sum 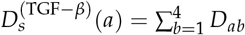 in decreasing order for *a* = 5, …, 44 (i.e. *a* belongs to one of the sets of up- or down-proteins) and where *D_ab_* is the sensitivity matrix for the 54 nodes of Table 1 (see also Figure 5).

Appendix Figure A4 provides an additional analysis of the overall influence of the TGF-*β* proteins on the up- and down-proteins which is discussed in subsection 3.4.

**Appendix Figure A5.**
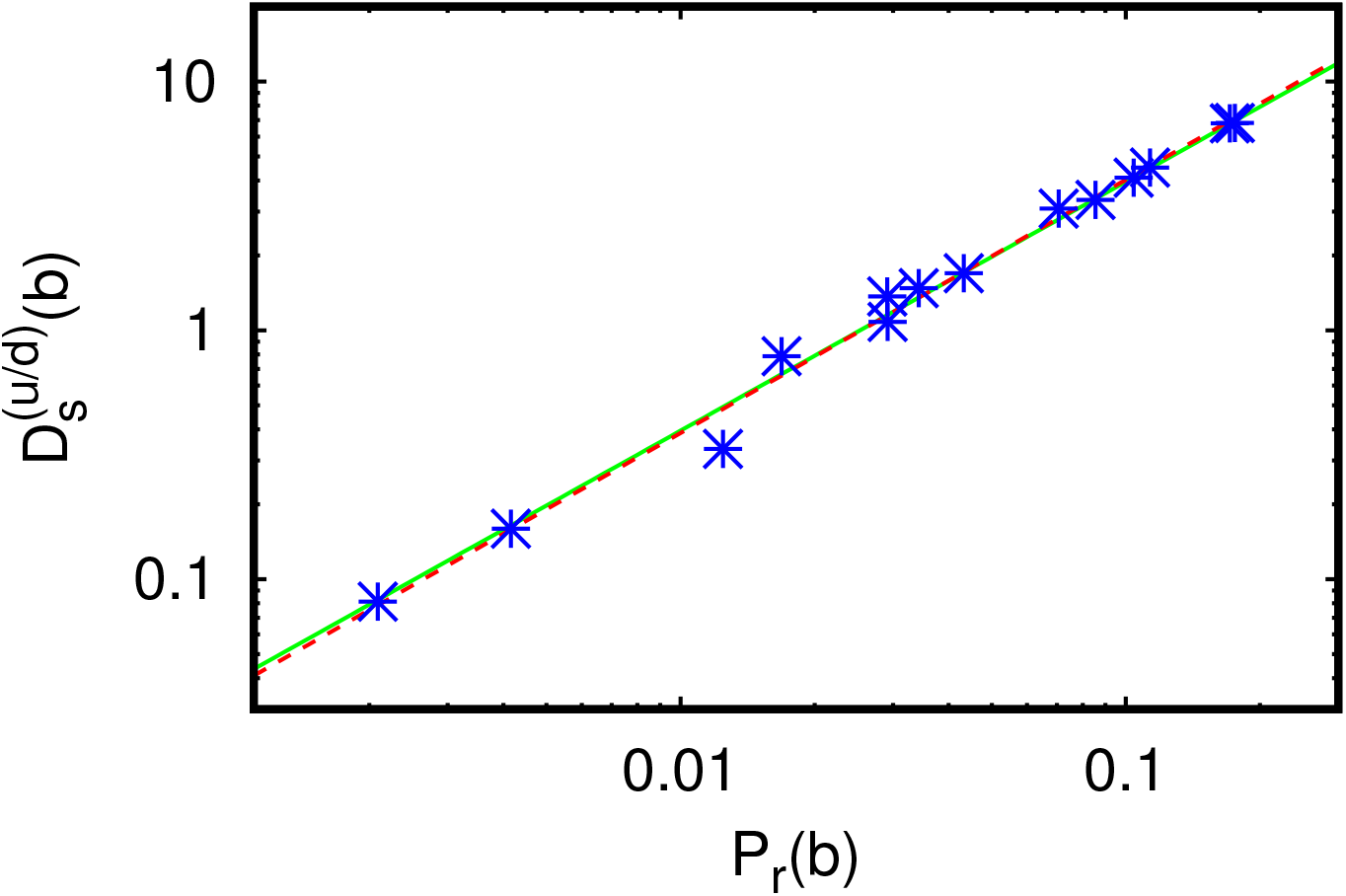
Dependence of the sum of sensitivities 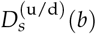 from Table 3 on the (local) PageRank probability *P*_r_ (*b*); the straight green line shows the fit dependence 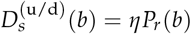 with the obtained numerical value *η* = 39.5 ± 1.4; the dashed red line corresponds to the power law fit 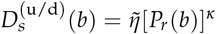 with 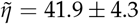 and *κ* = 1.017 ± 0.028.

Appendix Figure A5 provides the graphical and fit verification of the linear behavior between the two quantities 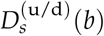 and *P*_r_(*b*) appearing the last two columns of Table 3.

### A2. Simple estimate for the sensitivity matrix

In the second part of this appendix we remind some details (see [29]) about the numerical computation of the sensitivity (8) and provide an analytic approximation based on a simplified model. Let (*a, b*) be an arbitrary index pair and *G_ε_* be the perturbed Google matrix obtained from a general unperturbed Google matrix *G*_0_ by multiplying its element *G*_0_(*a, b*) at position (*a, b*) by (1 + *ε*) and then sum-renormalizing the column *b* to unity. The elements in the other columns are not modified. In a more explicit formula we have:

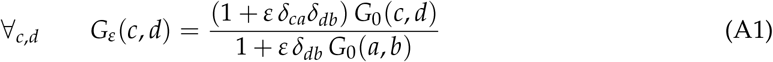

where *δ_ca_* = 1 (or 0) if *c* = *a* (or *c* ≠ *a*). Note that the denominator is either 1 if *d* ≠ *b* or the modified column sum 1 + *ε G*_0_(*a, b*) of column *b* if *d* = *b*. Expanding (A1) up to first order in *ε* we obtain *G_ε_* = *G*_0_ + *ε*Δ*G* + … with Δ*G* having the elements:

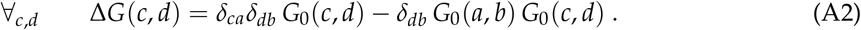

Let *P_ε_* be the sum-normalized PageRank vector of *G_ε_* determined by the conditions *G_ε_ P_ε_* = *P_ε_* and the normalization *E^T^ P_ε_* = 1 where *E^T^* = (1, …, 1) is a (row) vector with unit entries. Note that the column sum condition of *G_ε_* can be written as *E^T^ *G*_ε_* = *E^T^* and of course for *ε* = 0 we also have *G*_0_ *P*_0_ = *P*_0_, *E^T^ P*_0_ = 1 and *E^T^ G*_0_ = *E^T^*. Furthermore we write the perturbed PageRank vector in the form *P_ε_* = *P*_0_ + *ε*Δ*P* + … where the Δ*P* must satisfy the condition *E^T^* Δ*P* = 0. Then the sensitivity (8) is directly related to Δ*P* by:

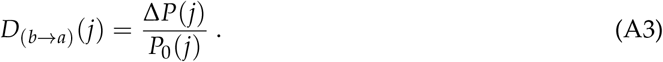

Expanding the PageRank equation *G_ε_ P*_ε_ = (*G*_0_ + *ε*Δ*G* + …)(*P*_0_ + *ε*Δ*P* + …) = *P_ε_* = *P*_0_ + *ε*Δ*P* + … to order one we first obtain the unperturbed PageRank equation *G*_0_ *P*_0_ = *P*_0_ and a further inhomogeneous equation :

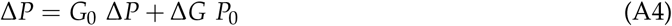

which can be efficiently numerically solved by iteration (choosing initially Δ*P* = 0 on the right hand side) once *P*_0_ has been computed (see [29] for details on this point). This provides a numerical precise scheme to compute the sensitivity in the limit *ε* → 0 without the need to take finite *ε*-differences.

Now, we consider a particular very simple model where *G*_0_ has identical columns being the PageRank *P*_0_, i.e. *G*_0_ = *P*_0_ *E^T^* or more explicitely *G*_0_ (*c*, *d*) = *P*_0_(*c*) for all values of *c, d*. Then we obtain from (A2)

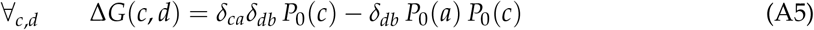

and from (A4)

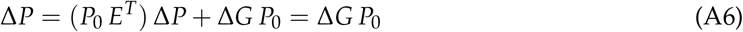

since *E^T^* Δ*P* = 0. Inserting (A5) in (A6) we obtain (replacing *c* = *j* and performing the *d*-sum for the matrix vector product)

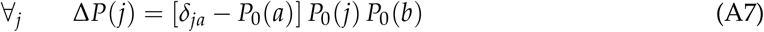

and from (A3)

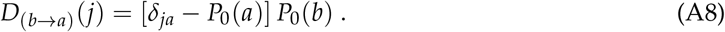

Choosing *j* = *a* this gives the sensitivity matrix

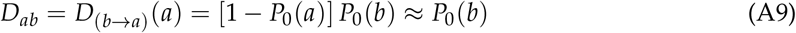

where the last approximation holds if typically *P*_0_(*a*) ≪ 1.

This result is of course only valid for the simplified model of identical columns (being the PageRank vector) in *G*_0_. However, when *G*_0_ represents a typical reduced Google matrix, with *N_r_* ≪ *N*, the component *G*_pr_ which has the strongest numerical weight (typically ~ 95%) is of the form 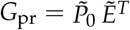 where 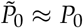 and 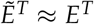 except for a few number of components *j* where strong deviations between 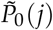 and *P*_0_(*j*) (and similarly between 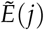 and *E*(*j*)) are possible.

Our examples of *D_ab_* visible in Figure 5 and of 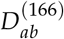 of Appendix Figure A3 confirm the typical behavior *D_ab_* ~ *P*_0_(*b*) for a “uniform background” but there are some exceptional peak values which arise from the deviations from *G*_pr_ to the simplified model and also from the contributions of *G*_rr_ and *G*_qr_. This also explains our numerical finding that all matrix elements of *D_ab_* are positive. Actually, according to (A8) we expect that *D*_(*b→a*)_ (*j*) is typically positive if *j* = *a* and negative if *j* ≠ *a*.

Furthermore, when taking the partial *a*-sum over up- and down-nodes of *D_ab_* the effect of exceptional peaks is strongly reduced thus explaining the linear behavior 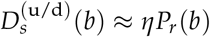 visible in Table 3 and Figure A5.

## Author Contributions

All authors equally contributed to all stages of this work.

